# Depth-discrete metagenomics reveals the roles of microbes in biogeochemical cycling in the tropical freshwater Lake Tanganyika

**DOI:** 10.1101/834861

**Authors:** Patricia Q. Tran, Samantha C. Bachand, Peter B. McIntyre, Benjamin M. Kraemer, Yvonne Vadeboncoeur, Ismael A. Kimirei, Rashid Tamatamah, Katherine D. McMahon, Karthik Anantharaman

## Abstract

Lake Tanganyika (LT) is the largest tropical freshwater lake, and the largest body of anoxic freshwater on Earth’s surface. LT’s mixed oxygenated surface waters float atop a permanently anoxic layer and host rich animal biodiversity. However, little is known about microorganisms inhabiting LT’s 1470 m deep water column and their contributions to nutrient cycling, which affect ecosystem-level function and productivity. Here, we applied genome-resolved metagenomics and environmental analyses to link specific taxa to key biogeochemical processes across a vertical depth gradient in LT. We reconstructed 523 unique metagenome-assembled genomes (MAGs) from 21 bacterial and archaeal phyla, including many rarely observed in freshwater lakes. We identified sharp contrasts in community composition and metabolic potential with an abundance of typical freshwater taxa in oxygenated mixed upper layers, and Archaea and uncultured Candidate Phyla in deep anoxic waters. Genomic capacity for nitrogen and sulfur cycling was abundant in MAGs recovered from anoxic waters, highlighting microbial contributions to the productive surface layers via recycling of upwelled nutrients, and greenhouse gases such as nitrous oxide. Overall, our study provides a blueprint for incorporation of aquatic microbial genomics in the representation of tropical freshwater lakes, especially in the context of ongoing climate change which is predicted to bring increased stratification and anoxia to freshwater lakes.

## Introduction

Located in the East African Rift Valley, Lake Tanganyika (LT) holds 16% of the Earth’s freshwater and is the second largest lake by volume. By its sheer size and magnitude, LT exerts a major influence on biogeochemical cycling on regional and global scales [1, 2]. For instance, LT stores over 23 Tg of methane below the oxycline [2], and about 14,000,000Tg of carbon in its sediments [1]. Over the past centuries, LT’s rich animal biodiversity has been a model for the study of species radiation and evolution [3]. In contrast, the microbial communities in LT that drive much of the ecosystem-scale productivity remains largely unknown.

LT is over 10 millions years old and oligotrophic, which provides a unique ecosystem to study microbial diversity and function in freshwater lakes, specifically tropical lakes. The comparatively thin, oxygenated surface layer of this ancient, deep lake harbors some of the most spectacular fish species diversity on Earth [4], but surprisingly ~80% of the 1890 km^3^ of water is anoxic. Being meromictic, its water column is permanently stratified. This causes a large volume of anoxic and nutrient-rich-bottom-waters to be thermally isolated from the upper ~70m of well-lit, nutrient-depleted surface waters. Despite stratification, periodically in response to sustained winds, pulses of phosphorus and nitrogen upwelling from deep waters replenish the oxygenated surface layers and sustain its productivity [5].

The physico-chemical environment of Lake Tanganyika is quite distinct from other ancient lakes[6–8]. For example, Lake Baikal, located in Siberia, is a seasonally ice-covered, of comparable depth (1642m for Lake Baikal, and 1470m for Lake Tanganyika), yet their thermal and oxygen profiles are drastically distinct. While Lake Tanganyika is a permanently stratified layer with a large layer of anoxic waters, Lake Baikal’s deepest layers are oxygenated.

Previous work on LT’s microbial ecology documented spatial heterogeneity in microbial community composition, especially with depth [9, 10]. The observed differences were primarily related to thermal stratification, which leads to strong gradients in oxygen and nutrient concentrations. Early evidence from LT suggests that anerobic microbially-driven nitrogen cycling such as anerobic ammonium oxidation (anammox) is an important component of nitrogen cycling [11]. However, the emergent effects of depth-specific variation in microbial communities on biogeochemical and nutrient cycling in LT remains largely unknown.

Here, we investigated microbial community composition, metabolic interactions, and microbial contributions to biogeochemical cycling along ecological gradients from high light, oxygenated surface waters to dark, oxygen-free and nutrient-rich bottom waters of LT. Our comprehensive analyses include genome-resolved metagenomics to reconstruct hundreds of bacterial and archaeal genomes which were used for metabolic reconstructions at the resolution of individual organisms and the entire microbial community, and across different layers in the water column. Additionally, we compared the diversity in two contrasting ancient, and deep rift-formed lakes (Lake Tanganyika and Lake Baikal) to address the question about ancient microbial lineages, evolution, and endemic lineages in freshwater lakes. Our work offers a window into the understudied microbial diversity of LT, provides a baseline for microbial lineages in deep lakes, serves as a case study for investigating microbial roles and links to biogeochemistry in globally distributed anoxic freshwater lakes.

## Methods

### Sample collection

Samples were collected in Lake Tanganyika, located in Central-East Africa, and based around the Kigoma and Mahale regions of Tanzania. We sampled near Kigoma because there are established field sites there and data that go back more than a decade. We sampled near Mahale National Park because of our focus on conservation in the nearly intact nearshore ecosystems there. Lake Tanganyika has a strong latitudinal gradient (>400 miles), therefore sampling in these two locations was also a way to capture some of that latitudinal variability.

From 2010 to 2013, samples for chlorophyll *a*, conductivity, DO, and temperature were collected. Secchi depths were collected over two years in 2012 and 2013, from a previous research cruise using a YSI 6600 sonde with optical DO and chlorophyll-a sensors. Those measurements were collected down to approximately 150m. Twenty-four water samples were collected in 2015 for metagenome-sequencing. A summary of all samples are available in **Figure S1**. The samples were collected around two stations termed Kigoma and Mahale based on nearby cities. Water samples were collected with a vertically oriented Van Dorn bottle in 2015 and metadata is listed in **Table S1**. For the metagenomes, three Van Dorn bottle casts (10L) in Kigoma and Mahale were collected, in which depth-discrete samples were collected across a vertical gradient, down to 1200m at the maximum depth (**Figure 1**, **Table S1**). Additionally, five surface samples were collected from the Mahale region, and one surface sample from the Kigoma region (**Figure 1**). Methods to produce the map in **Figure 1** using the geographic information system (GIS) ArcMap are described in the **Supplementary Text.**

**Figure 1.**
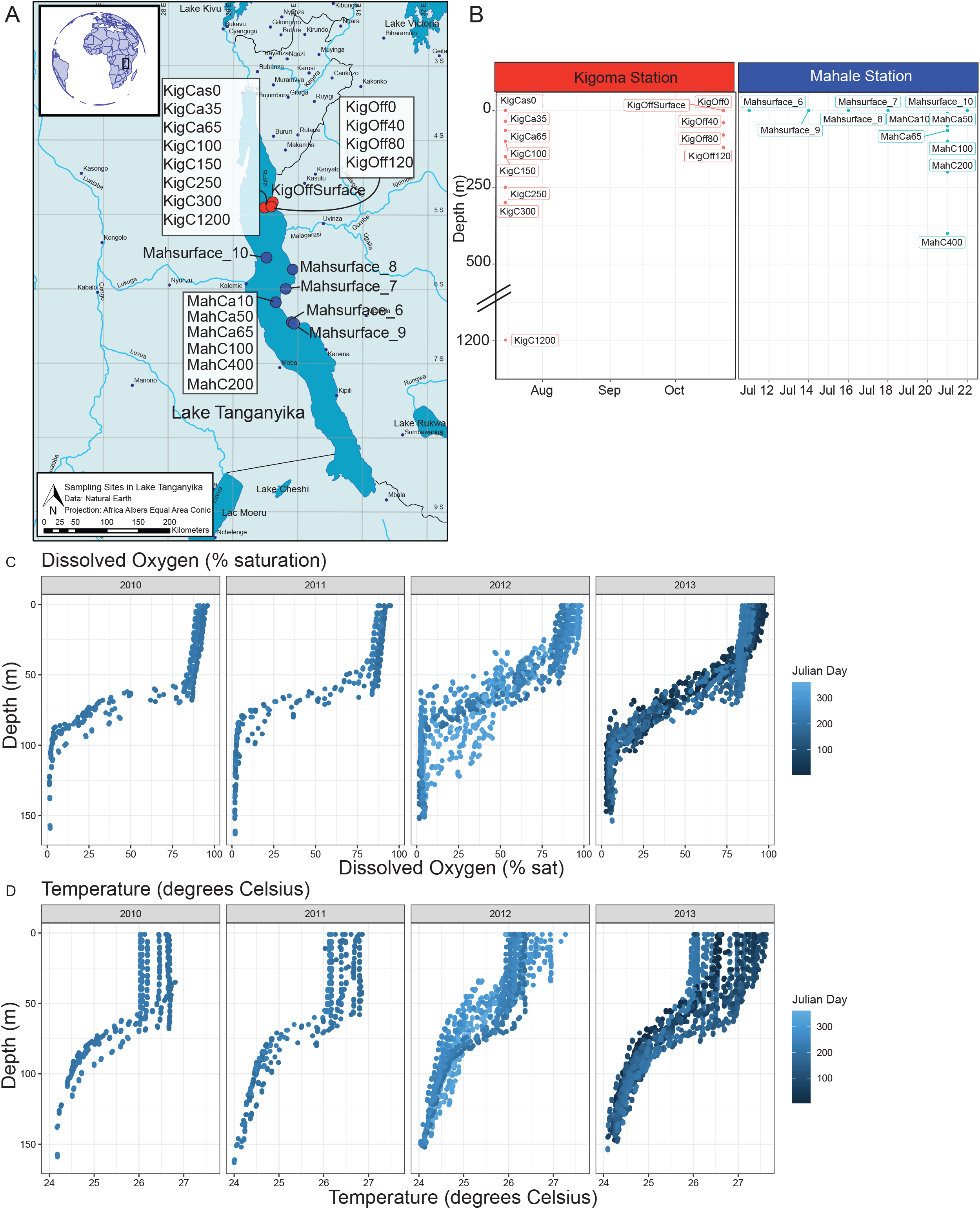
**A.** Map of sampling locations in LT. The inset figure shows the approximate location of LT in a spherical global projection. Lakes and rivers are identified in blue; cities are shown in black. Rivers, lakes, and populous cities are labeled. The locations of the 24 samples collected are labelled, corresponding to Kigoma(red) or Mahale (blue), which are stations in Lake Tanganyika. The white shaded box represents location for which we collected depth-discrete samples (cast). **B.** Sampling locations along the vertical transect of LT. Samples are depicted foremost by location (Kigoma vs. Mahale), followed by sampling date on the x-axis and depth on the y-axis. Short sample names are written next to each dot. **C.** Dissolved oxygen (% saturation) in Lake Tanganyika, in percentage of saturation, from 2010 to 2013. **D.** Temperature profiles, in degree Celsius, from 2010 to 2013 in Lake Tanganyika.

We filtered as much as we could until the filter clogged, which was usually between 500-1000 mL. This usually took about 10-20 min of hand pumping. The water was not prefiltered, but filtered directly through a 47mm 0.2 um pore nitrocellulose filter which was stored in a 2 mL tube with RNAlater. Filters were frozen immediately and brought back on dry ice to UW-Madison.

### DNA Extraction and Sequencing

DNA extractions were performed using the MP Biomedicals FastDNA Spin Kit with minor protocol modifications as described previously [12] back in Madison, WI. Metagenomic DNA was sequenced at the Joint Genome Institute (JGI) (Walnut Creek, CA) on the Illumina HiSeq 2500 platform (Illumina, San Diego, CA, U.S.A.), which produces 2 x 150 base pairs (bp) reads with a targeted insert size of ~240 bp.

### Metagenome Assembly, Genome Binning, Dereplication and Selection of Genomes

Each of the 24 individual samples were assembled *de novo* by JGI to obtain 24 metagenomes assemblies. Briefly, raw metagenomic sequence reads were quality filtered, then assembled using MetaSPADEs v3.12 (default parameters) [13]. Each metagenome was binned individually using: MetaBat1 [14], MetaBat2 [15] v2.1.12 and MaxBin2 v2.2.4 [16]. We consolidated the bins generated by these three different tools using DASTool v1.1.0 [17] (default parameters except for --score_threshold 0.4), resulting in a total of 3948 MAGs. We dereplicated these MAGs using dRep v2.3.2 with the default settings[18]. A total of 821 unique clusters were created. Genome completion was estimated using CheckM v1.0.11 [19] and DAStool. To select a set of medium and high-quality genomes (≥50% completeness, <10% contamination) for downstream analyses, we used the minimum genome information standards for metagenome-assembled genomes [20] which yielded a total of unique dereplicated 523 MAGs (**Table S2**). In total, these representative MAGs represent clusters containing 3948 MAGs. The 523 MAGs are used as the dataset for this study.

### Relative abundance across water column depths and Gene Annotations

We mapped each metagenomic paired-read set to each of the 523 MAGs using BBMap [21], with default settings, to obtain a matrix of relative coverage (as a proxy for abundance across the samples) versus MAG. We used pileup.sh implemented in BBMap which calculates the coverage values, while also normalizing for the length of the scaffold and genome size[21]. We combined the mapping table from the 24 metagenomes, and summed the total coverage based on the associated MAGs identifier for each scaffold. To normalize the coverage by the metagenome size, we divided the coverage by the number of total reads per metagenome. Open reading frames (ORFs) of the scaffolds were identified using Prodigal v2.6.3 [22].

### Identification of phylogenetic markers

A set of curated Hidden Markov Models (HMM) for 16 single-copy ribosomal proteins (rpL2, rpL3, rpL4, rpL5, rpL6, rpL14, rpL14, rpL15, rpL16, rpL18, rpL22, rpL24, rpS3, rpS3, rpS8, rpS10, rpS17, rpS19) [23] was used to identify these genes in each MAG using hmmsearch (HMMER 3.1b2) [24] with the setting --cut_tc (using custom derived trusted cutoffs). The esl-reformat.sh from hmmsearch was used to extract alignment hits.

### Phylogenetic tree

To create the concatenated gene phylogeny, we used reference MAGs which represented a wide range of environments including marine, soil, hydrothermal environments, coastal and estuarine environments originally built from [25, 26]. We used hmmsearch to identify the16 ribosomal proteins for the bacterial tree, and 14 ribosomal proteins for the archaeal tree as described previously[25] (also described in the section “Identification of phylogenetic markers”). All identified ribosomal proteins for the backbone and the LT MAGs were imported to Geneious Prime V.2019.0.04 (https://www.geneious.com) separately for Bacteria and Archaea. For each ribosomal protein, we aligned the sequences using MAFFT (v7.388, with parameters: Automatic algorithm, BLOSUM62 scoring matrix, 1.53 gap open penalty, 0.123 offset value) [27]. Alignments were manually verified: in the case that more than one copy of the ribosomal protein was identified, we performed a sequence alignment of that protein using MAFFT (same settings) and compared the alignments for those copies. For example, if they corresponded exactly to a split protein, we concatenated them to obtain a full-length protein. If they were the same section (overlap) of the protein, but one was shorter than the other, the longer copy was retained. We applied a 50% gap masking threshold and concatenated the 16 (or 14) proteins. The concatenated alignment was exported into the fasta format and used as an input for RAxML-HPC, using the CIPRES server [28], with the following settings: datatype = protein, maximum-likelihood search = TRUE, no bfgs = FALSE, print br length = false, protein matrix spec: JTT, runtime=168 hours, use bootstrapping = TRUE. The resulting Newick format tree was visualized with FigTree (http://tree.bio.ed.ac.uk/software/figtree/). The same procedure was followed to create the taxon-specific tree, for example for the Candidatus Tanganyikabacteria and Nitrospira, using phylum-specific references except with no gap masking since sequences were highly similar and had few gaps. Additionally, an *amoA* gene phylogeny was performed using UniProt *amoA* sequences with the two “predicted” comammox genomes with 90% masking. (**Supplementary Text**).

### Taxonomic assignment and comparison of manual versus automated methods

Taxonomic classification of MAGs was performed manually by careful inspection of the RP16 gene phylogeny, bootstrap values of each group, and closest named representatives (**Supplementary Material 1-2**). Additionally, we also assigned taxonomy using GTDB-tk [29], which uses ANI comparisons to reference genomes and 120 marker genes, using FastTree (**Supplementary Material 4**). We compared the results for taxonomic classification between the manually curated and automated approaches to check whether taxonomic assignment matched across phyla, and finer levels of resolution. Since the two trees (RP16 and gtdb-tk tree) are made using different sets of reference genomes, we are unable to do a direct comparison of the taxonomic position of each of the MAGs in our study. Therefore, we manually curated each manual vs. GTDB-tk classification and added a column stating whether the results matched (**Table S2)**.

To enhance the MAGs relevance to freshwater microbial ecologists, we assigned taxonomic identities to 16S rRNA genes identified in our MAGs to names in the guide on freshwater microbial taxonomy [30]. 16S rRNA genes were identified in 313 out of 523 MAGs using CheckM’s ssu_finder function[19]. The 16S rRNA genes were then used as an input in TaxAss[31], which is a tool to assign taxonomy to the identified 16S rRNA sequences against a freshwater-specific database (FreshTrain). This freshwater-specific database, albeit focused on epilimnia of temperate lakes, is useful for comparable terminology between the LT genomes and the “typical” freshwater bacterial clades, lineages and tribes terminology defined previously [30]. We also compared taxonomic identification among the three methods and show the results in **Table S2.**

We chose to provide the information from all sources of evidence (RP16, GTDB-tk tree, manual curation, automated taxonomic classification and 16S rRNA sequences), even if in instances some results might be inconsistent. It is important to note that each source of evidence and reference-based methods are biased by their database content, but we hope that providing several lines of evidence can help arrive to a consensus. For readability, the MAGs in our study are referred by their manually curated phylum or lineage name.

### Support for Tanganyikabacteria, a monophyletic sister lineage to Sericytochromatia

We noted 3 MAGs from Lake Tanganyika to be monophyletic, but initially were unrelated to any reference genomes. Since they were placed close to the Cyanobacteria, we created a detailed RP16 tree of 309 genomes of Cyanobacteria, and sister-lineages (Sericytochromatia/Melainabacteria [32], Blackallbacteria, WOR-1/Saganbacteria, and Margulisbacteria) including other freshwater lakes [33, 34], groundwater marine, sediment, fecal, isolates and other environments. We calculated pairwise genome-level average nucleotide identities (ANI) values between all 309 vs 309 genomes using fastANI [35]. The 3 genomes from Lake Tanganyika were closely related, yet distinct from, the recently defined Cyanobacterial class Sericytochromatia[32, 36], using evidence from RP16 and GTDB-tk phylogeny of the MAGs, and average nucleotide identity (ANI) with closely related genomes in the literature (**Table S3**). The other existing Sericytochromatia used for comparison were isolated from Rifle acetate amendment columns (Candidatus Sericytochromatia bacterium S15B-MN24 RAAC_196; GCA 002083785.1) and coal bed methane well (Candidatus Sericytochromatia bacterium S15B-MN24 CBMW_12; GCA 002083825)

### Comparison of MAG taxonomic diversity in Lake Tanganyika versus Lake Baikal

To compare taxonomic diversity in two ancient deep lakes, Lake Tanganyika and Lake Baikal, we compare the ANI of metagenome-assembled genomes from the deepest samples from the two lakes. Lake Baikal has 231 MAGs published[8]. These 231 MAGs were assembleld from samples from depths of 1350 and 1250m. We selected all MAGs that had a read abundance ≥ 0.5% of the KigC1200 sample (1200m depth) microbial diversity, resulting in 260 MAGs. We used fastANI to compare all vs. all (491 vs. 491 MAGs) %ANI values. To assess the patterns, we generated histograms and mean ANI values and plotted them in R. We grouped the pairwise matches as Tanganyika vs. Tanganyika, Baikal vs. Baikal, and Tanganyika vs. Baikal (which is the same as Baikal vs. Tanganyika).

### Metabolic potential analysis and comparison of metabolic potential and connection across three distinct depths in LT

Metabolic potential of LT MAGs was assessed using METABOLIC which includes 143 custom HMM profiles [37], using hmmsearch (HMMER 3.1b2) (--use_tc option) and esl-reformat to export the alignments for the HMM hits [24]. We classified the number of genes involved in metabolism of sulfur, hydrogen, methane, nitrogen, oxygen, C1-compounds, carbon monoxide, carbon dioxide (carbon fixation), organic nitrogen (urea), halogenated compounds, arsenic, selenium, nitriles, and metals. To determine if an organism could perform a metabolic function, one copy of each representative gene of the pathway must have been present in the MAG, for which a value of 1 (presence) was written, as opposed to 0 (absence). To investigate heterotrophy associated with utilization of complex carbohydrates, carbohydrate degrading enzymes were annotated using hmmscan on the dbCAN2[38] (dbCAN-HMMdb-V7 downloaded June 2019) database.

The sample profile collected on July 25, 2015 covered a vertical gradient ranging from surface samples to 1200m. We analyzed samples collected near Kigoma from 3 different depths (0m (KigCas0), 150 m (KigCas150), 1200m (KigCas1200)) based on the general difference in abundance of key taxa. We used METABOLIC v4.0 [37] to identify and visualize organisms with genes involved in carbon, sulfur, or nitrogen metabolism. We combined the results into a single figure showing the number of genomes potentially involved in each reaction, and the relative community relative abundance of those organisms across 3 distinct depths. We performed some additional manual curation for differentiating between *amo* and *pmo* genes, and annotating *nxr* genes (**Supplementary Text**).

In addition to the functional gene annotations done by METABOLIC, which includes HMMs related to biogeochemical cycles, all 24 metagenomic assemblies were annotated by IMG/M, pipeline version 4.15.1[39].

## Results

We sampled a range of physico-chemical measurements in LT which are described in **Figure 1** and **Figure S1**. We collected 24 metagenome samples from the LT water column spanning 0 to 1200m, at two stations Kigoma and Mahale in 2015 (**Figure 1B**), which consisted of 18 depth-discrete samples and six were surface samples. Despite not having paired physiochemical profiles in 2015, temperature and dissolved oxygen from 2010 to 2013 was highly consistent and showed minimal interannual variability, particularly during our sampling period (July and October) (**Figure 1**, **Figure S2**). Additionally, the environmental profiles are similar to those collected in other studies [40–43]. Water column temperatures ranged from 24 to 28°C, and changes in DO were greatest at depths ranging from ~50 to 100 m during the time of sampling dropping to 0% saturation DO around 100m. The thermocline depths shift vertically depending on the year (**Figure S2**). Secchi depth, a measure of how deep light penetrates through the water column was on average 12.2m in July and 12.0m in October (**Figure S3**). Nitrate concentrations increased up to ~100 μg/L at 100m deep, followed by rapid depletion with the onset of anoxia (**Figure S4**). A consistent chlorophyll-a peak was detected at a depth of ~120m in 2010-2013 (**Figure S5**). Comparatively in 2018, the peak occurred around 50m[43]. Other profiles collected in 2018 show a nitrate peak between 50m and 100m in LT[43].

### The microbiome of Lake Tanganyika

Metagenomic sequencing, assembly, and binning resulted in 3948 draft-quality MAGs that were dereplicated and quality-checked into a set of 523 non-redundant medium-to high-quality MAGs for downstream analyses. To assign taxonomic classifications to the organisms represented by the genomes, we combined two complimentary genome-based phylogenetic approaches: a manual RP16 approach, and GTDB-tk [29], an automated program which uses 120 concatenated protein coding genes. For the most part, we observed congruence between the two approaches (**Figure S6**) and the genomes represented 24 Archaea and 499 Bacteria from 21 phyla, 23 classes, and 38 orders (**Table S2, Figure 2**). While manually curating the RP16 tree, and upon closer inspection and phylogenetic analysis of Cyanobacterial genomes and sister lineage genomes (Sericytochromatia, Melainabacteria, etc.), we noticed that three bacterial genomes formed a monophyletic freshwater-only clade within Sericytochromatia, sharing less than 75% genome-level average nucleotide identity (ANI) to other non-photosynthetic Cyanobacteria-like sequences (**Figure 3, Table S3**). On this basis, we propose to name this lineage Candidatus Tanganyikabacteria (named after the lake). Sericytochromatia is a class of non-photosynthetic Cyanobacteria-related organisms, that has recently gained attention along with related lineages such as Melainabacteria and Margulisbacteria due to the lack of photosynthesis genes, indicating phototrophy was not an ancestral feature of the Cyanobacteria phylum [32, 36]. This is the first recovery of Sericytochromatia from freshwater lake environments, since others were found in glacial surface ice, biofilm from a bioreactor, a coal bed methane well, and an acetate amendment column from the terrestrial subsurface.

**Figure 2.**
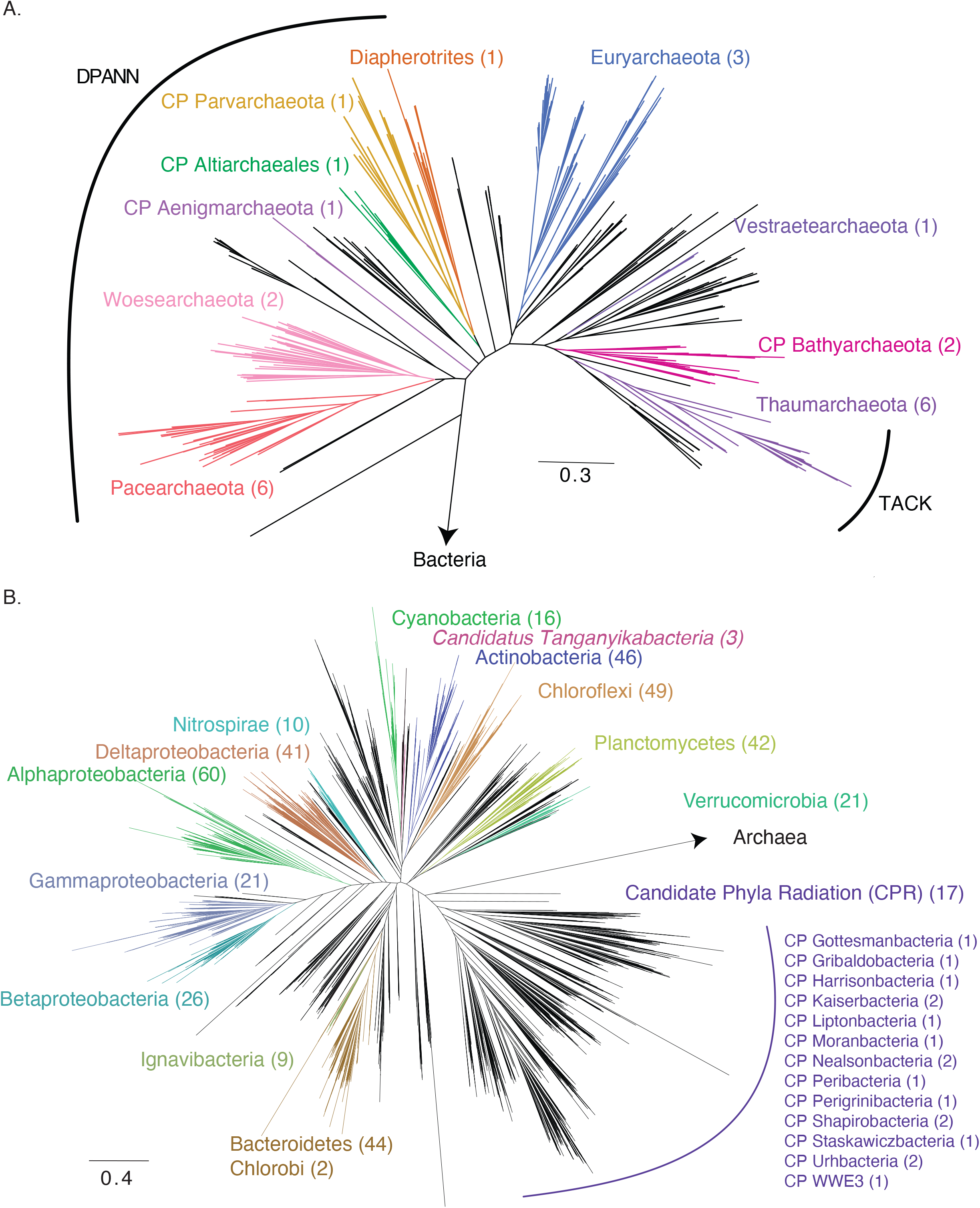
Phylogeny of (A) Archaeal and (B) Bacterial metagenome-assembled genomes (MAG) recovered from LT. The tree was constructed using 14 and 16 concatenated ribosomal proteins respectively, and visualized using FigTree. Not all lineages are named on the figure. The number of medium and high quality MAGs belonging to each group are listed in parentheses. Colored groups represent the most abundant lineages in LT. Tanganyikabacteria MAGs from this study are italicized. A more detailed version of the tree constructed in iTol is found in Supplementary Files 1 and 2 with bootstrap values and names of taxa.

**Figure 3.**
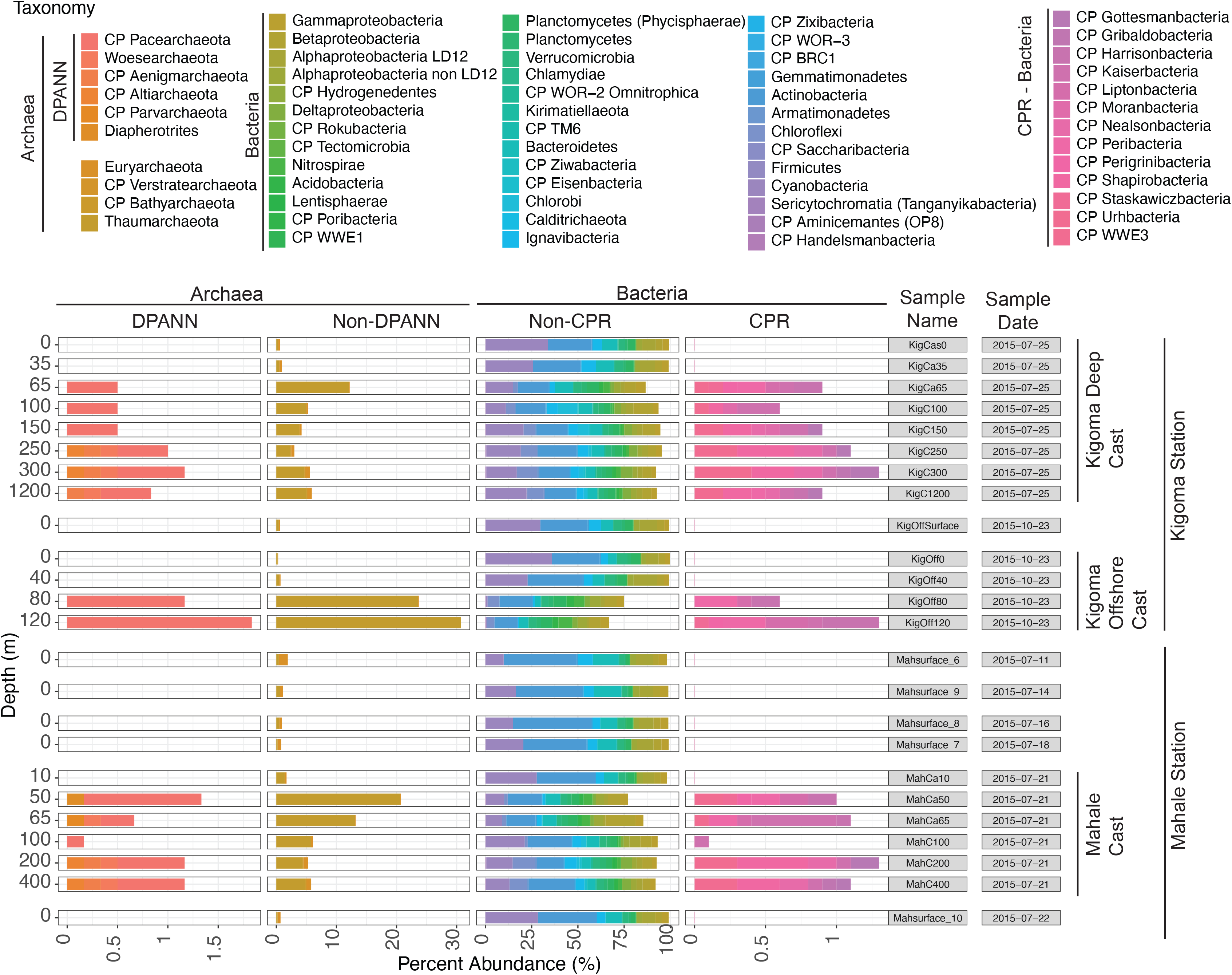
**A.** Concatenated genome phylogeny using 16 ribosomal proteins of metagenome-assembled genomes of Cyanobacteria and sister-lineages of non-photosynthetic Cyanobacteria such as Margulisbacteria (WOR-1), Melainabacteria and Sericytochromatia. The 3 MAGs from Lake Tanganyika are labeled in blue, and represent a high-support (100) monophyletic lineage among the known Sericytochromatia. The presence-absence plot shows gene involved in oxygen metabolism (squares) and nitrogen metabolism (circles). The MAG from Lake Tanganyika is the only one among all genomes to have genes for denitrification. M_DeepCast_65m_m2_071 has 96.58% genome completeness (Full un-collapsed tree available in Supplementary Material). **B.** Genome-level average nucleotide identity (ANI) (%) are shown as pairwise matrix for the Sericytochromatia 5 MAGs boxed.

To investigate the stratification of microbial populations and metabolic processes in the water column of LT, we identified three zones based on oxygen saturation. The microbial community composition and community relative abundance (calculated as relative abundance of reads mapped, RAR) across our samples (**Figure S7**) was distinct between the oxygenated upper layers and the deep anoxic layers (**Figure 4, Table S4)**. Notably, Archaea accounted for up to 30% of RAR in sub-oxic samples (Kigoma 80 and 120m), and generally increased in abundance with depth. DPANN archaea were only found in sub-oxic (>50m deep) samples. Candidate phyla radiation (CPR) organisms generally increased in abundance with depth, reaching up to ~2.5% RAR. Among bacteria, notably, common freshwater taxa such as Actinobacteria (e.g. acI, acIV) Alphaproteobacteria (LD12), and Cyanobacteria showed a ubiquitous distribution, whereas certain groups such as Chlorobi and Thaumarchaetota were most abundant below 100m. The community structure composition and abundance of the surface samples in the five Mahale samples was similar throughout. We note that some MAGs were recovered from samples (included in the name of the MAG) in which they had comparatively low RAR but were much more abundant in other samples.

**Figure 4.**
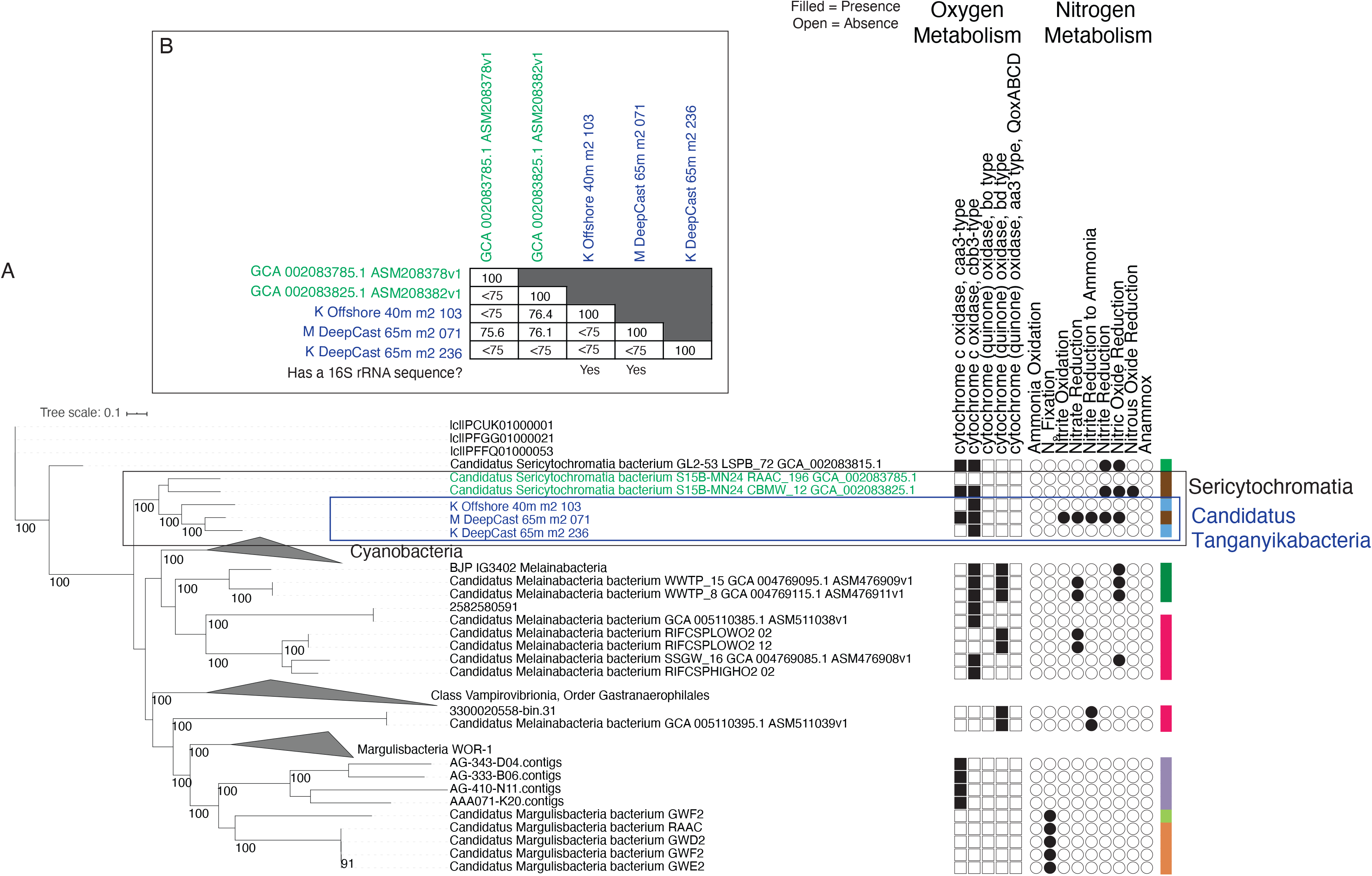
Bar plot showing the relative read abundance (RAR) per sample in the 24 metagenome from Lake Tanganyika. MAGs are taxonomically ordered along the x-axis, with closely related taxa next to each other. Additionally, information about whether they are Archaea (DPANN or not) and Bacteria (CPR or not) is specified. The three casts are identified with a vertical line on the right hand side. Samples are organized by location, then sampling date, then depth.

### How similar is the Lake Tanganyika microbiome to that of other deep and ancient freshwater lakes?

While a comparison of Lake Tanganyika’s metagenomic diversity would be interesting to compare with other African Great Lakes (such as Lake Malawi, Lake Kivu, Lake Victoria), no published MAGs from those sites existed at the time of writing. We compared Lake Tanganyika and Lake Baikal’s (LB’s) microbial diversity (based on MAGs) because they are both ancient, extremely deep lakes, and therefore might both have sufficient evolutionary time for a wide diversity of bacterial and archaeal lineages to evolve. Yet, both lakes differ drastically in terms of environmental settings (LB is seasonally ice-covered with a oxygenated hypolimnion, whereas LT is a tropical ice-free lake with an anoxic bottom water layer). To compare the microbial communities, we compared the MAGs recovered from LT and LB (**Figure 5, Table S5, Figure S8)**.

**Figure 5.**
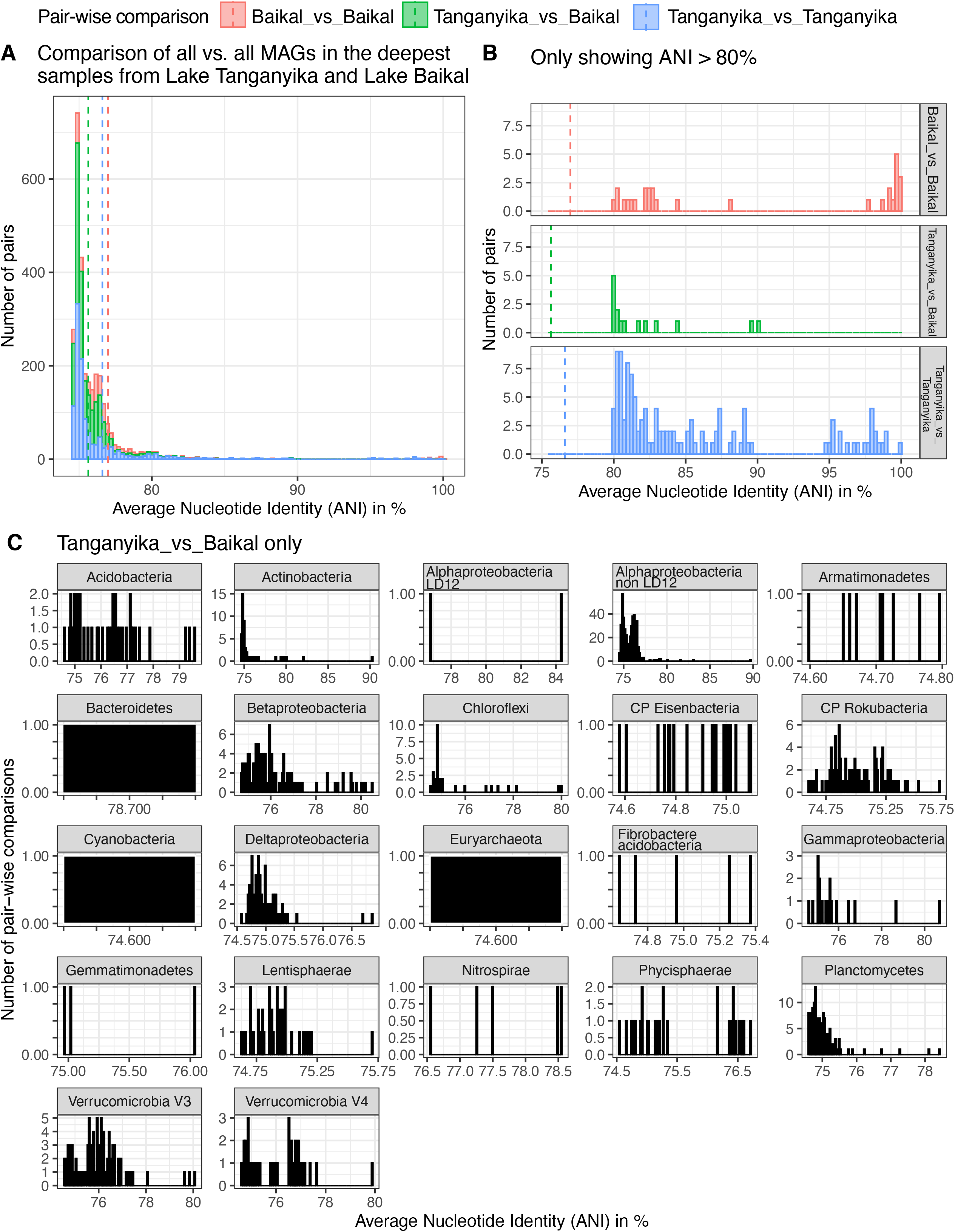
**A.** Histogram of the ANI values when doing pairwise comparison of full-genome MAGs from Lake Tanganyika vs. Lake Baikal. The vertical dashed lines correspond to the mean value (Baikal vs. Baikal: 79.98%, Tanganyika vs. Baikal: 75.63%, and Tanganyika vs. Tanganyika 76.60%). **B.** Zoom in ANI values above 80% only. There are slightly more ANI above the 97% ANI for Baikal vs. Baikal, than Tanganyika vs. Tanganyika. Two genomes from Baikal had 100% ANI, two genomes from Tanganyika (K_Offshore_surface_m2_005 and K_DeepCast_100m_m2_150.fasta) shared 99.91 %ANI. The highest %ANI between a MAG from Tanganyika vs. Baikal was among M_surface_10_m2_136.fasta and GCA_009694405.1_ASM969440v1_genomic.fna (90.19 %ANI) which were Candidatus Nanopelagicaceae bacterium, an Actinobacteria. **C.** Of the pairs in “Tanganyika_vs_Baikal” category, now colored by taxonomic groups, showing the number of pair-wise comparisons for each %ANI value, organized by taxonomic group. For example, there are only 1 pairs of Bacteroidetes, Cyanobacteria and Euryarchaeota from the two lakes that have ANI >75%.

Comparing the MAGs from the 1200m sample in LT (KigOff1200m, >0.05% RAR) and MAGs from deep samples (1200 and 1350m) from Baikal, relatively high RAR of Thaumarchaeota were identified in both lakes. Thaumarchaeaota accounted for ~10% RAR in LT and ~20% in LB. Typical freshwater taxa such as the acI lineage of Actinobacteria and LD12 group of Alphaproteobacteria (Ca. *Fonsibacter))* were observed in higher abundance in surface samples but also at depths up to 50m. CPR bacteria were found in both lakes. Archaea accounted for 6.3% of RAR in KigOff1200m, and about 2% in Lake Baikal. Archaeal MAGs from KigOff1200 (and >0.05% RAR) belonged to the lineages Bathyarchaeota, Verstraetaerchaeota, Thaumarchaeota, Euryarchaeota and DPANN (Pacearchaeota, Woesearchaeota, Aenigmarchaeota). The 3 DPANN MAGs from LB were closely related to Pacearchaeota and Woesearchaeota.

Interestingly, the majority of taxonomic groups observed in the deepest waters of LT (1200 m) were unique to LT (59%), whereas 68% of taxa richness recovered from LB that were bacterial MAGs were also identified in the deepest LT sample. Nitrososphaerales (Archaea) were identified in both LT and LB. Both LT and LB have a small abundance of Cyanobacteria. In LB, they account for ~1%, and are likely sourced from vertical mixing or sediment. However in LT, Cyanobacteria are significantly more abundant, accounting for 14 %RAR, but the lake is stratified with no mixing. Organisms from the lineage Desulfobacterota, with prominent roles in sulfur cycling (H2S generation) were identified in LT but not in LB. Overall, both lakes had high abundance and richness of Archaea and CPR, although the specific linneages (for example Desulfubacterota) likely reflect the differences in geochemistry in the lake, and to the different oxygen niches that these organisms may occupy.

### Depth-dependent contrasts in microbial metabolism in LT: Biogeochemical cycling of Carbon, Nitrogen and Sulfur

We investigated microbial metabolic potential and process-level linkages between MAGs in the surface, at 150m, and at 1200m, representing three distinct ecological layers within the lake (**Figure 6, Table S6-7**). Individual metabolic pathways were identified in each MAG and their capacity to contribute to carbon (**Figure S9, Table S6-7**), nitrogen (**Figure S10, Table S6-7**), and sulfur (**Figure S11, Table S6-7**) and metal biogeochemical cycles were assessed (**Figure S12, Table S6-7**). Based on historical environmental data[40], light sufficient to support photosynthesis is available to ~70m, and these surface waters are oxygen rich and nutrient depleted. The 150m depth is subject to more interannual year variation in temperature and DO (**Figures S2-5**). The 1200m sample represents a relatively stable, dark, nutrient-rich, and anoxic environment, according to data collected from 2010-2013 in our study and in accordance with historical profiles.

**Figure 6.**
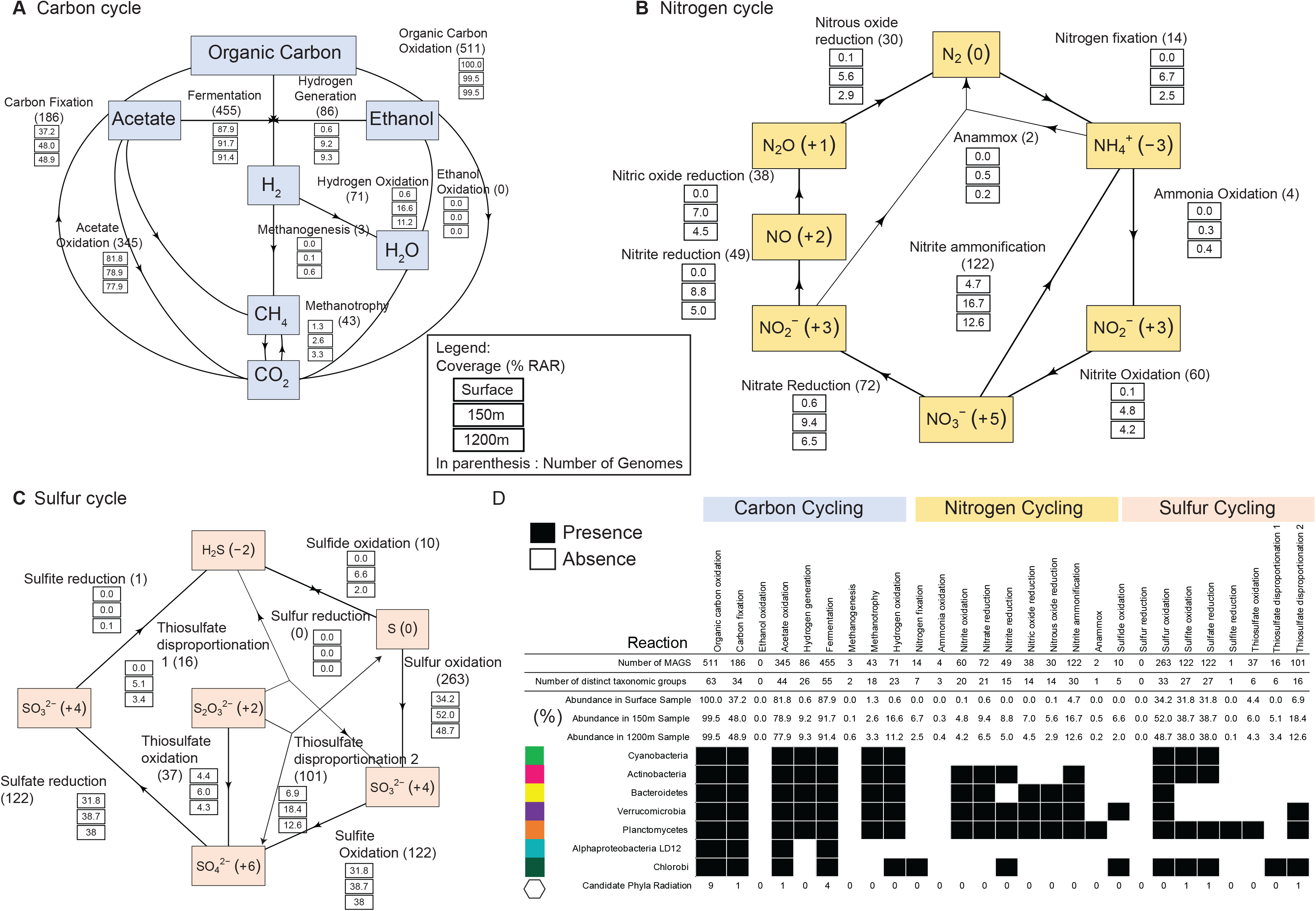
Abundance of organisms involved in different (**A**) carbon, (**B**) nitrogen and (**C**) sulfur cycling steps, in three depth-discrete samples from Kigoma (surface, 150m, 1200m).Oxidation states are shown in parentheses. **D.** Presence and absence of individual steps in C, N, S cycling. Abundance of organisms are described in percentages. Only reactions in a subset of common freshwater taxa (Cyanobacteria, Actinobacteria, Bacteroidetes, Verrucomicrobia, Planctomycetes, Alphaproteobacteria (LD12), Chlorobi) and Candidate Phyla Radiation are shown. Presence and absence of these pathways in taxonomic groups are represented by filled and open circles respectively.

Microbial pathways associated with the carbon cycle included the use of organic carbon (e.g. sugars), ethanol, acetate, methanogenesis, methanotrophy, and CO2-fixation (**Figure 6**). As would be expected, organisms capable of organic carbon oxidation were abundant comprising nearly 99% of the relative abundance of reads (RAR) community in the sub-oxic and anoxic samples. The metabolic potential for fermentation and hydrogen generation was also abundant in the sub-oxic and anoxic samples, represented by 91% RAR (455 MAGs) and 9% RAR (86 MAGs), respectively. Other anerobic processes including methanogenesis (3 MAGs) and methanotrophy (43 MAGs) were identified in a limited number of MAGs, but were observed to be more abundant in the 1200m samples. Both bacteria and archaea encoded carbohydrate-degrading enzymes (CAZYmes) (**Figure S13, Table S8**). The highest densities of CAZYmes (when normalized by genome size) were identified organisms from the lineages Verrucomicrobia, Lentispheara, and CP Shapirobacteria. The highest densities of CAZYmes (when normalized by genome size) were identified in organisms from the lineages Verrucomicrobia, Lentisphaerae, and CP Shapirobacteria. Meanwhile, Glycoside hydrolases (GHs) that participate in the breakdown of different complex carbohydrates were prominent in Verrucomicrobia and Planctomycetes, as has been observed in other studies of these common freshwater linneages in lakes[44, 45].

We identified microorganisms involved in the inorganic nitrogen cycle, including oxidation and reduction processes (**Figure 6, Table S6-7**). The metabolic potential for nitrogen cycling was generally higher in the sub-oxic samples as compared to the anoxic samples. Nitrogen fixation capacity (14 MAGs) was found across all 3 depths, and nitrogen fixing organisms comprised 0-6% RAR at each depth. We identified a MAG belonging to the Thaumarchaeal class Nitrososphaeria that contains Archaeal *amoABC* genes for ammonia oxidation (**Figure S14**), and many non-CPR Bacteria with *nxr* genes involved in nitrite oxidation (**Figure S15**). Nitrospira bacteria were identified to be capable of complete ammonia oxidation (comammox) (Clade II-A), with highest RAR in the sub-oxic depths (**Figure S16**). Denitrification processes that remove nitrate from the system were discovered across all depths, at approximately the same RAR in all depths. For example, the potential for nitrate reduction was identified in 72 MAGs that comprised 6.5-9% RAR and for nitrite reduction in 49 MAGs, at 5-9% RAR in anoxic depths. The nitrate/nitrite ammonification (DNRA) potential was identified in 122 MAGs accounting for between 5-17% RAR in each depth. Bacteria capable of anammox accounted for between 0.2-0.5% RAR at each depth.

Sulfur biogeochemical profiles exist in LT and generally show an increase in H2S beginning in the sub-oxic zone into the anoxic zone with the highest concentration observed in the deepest waters [40]. H2S can accumulate in the water column through sulfite reduction, thiosulfate disproportionation thiosulfate disproportionation into H2S and SO_3_^2-^, and sulfur reduction. While only a few MAGs were potentially able to perform these processes (1, 16 and 10 respectively), we observed that all of them increased in RAR with depth and were more abundant in the anoxic depths (**Figure 6, Figure 7**). Sulfur oxidation (263 MAGs), sulfite oxidation (122 MAGs), and sulfate reduction were each prominently represented processes, with such organisms accounting for over 30% RAR at each depth.

**Figure 7.**
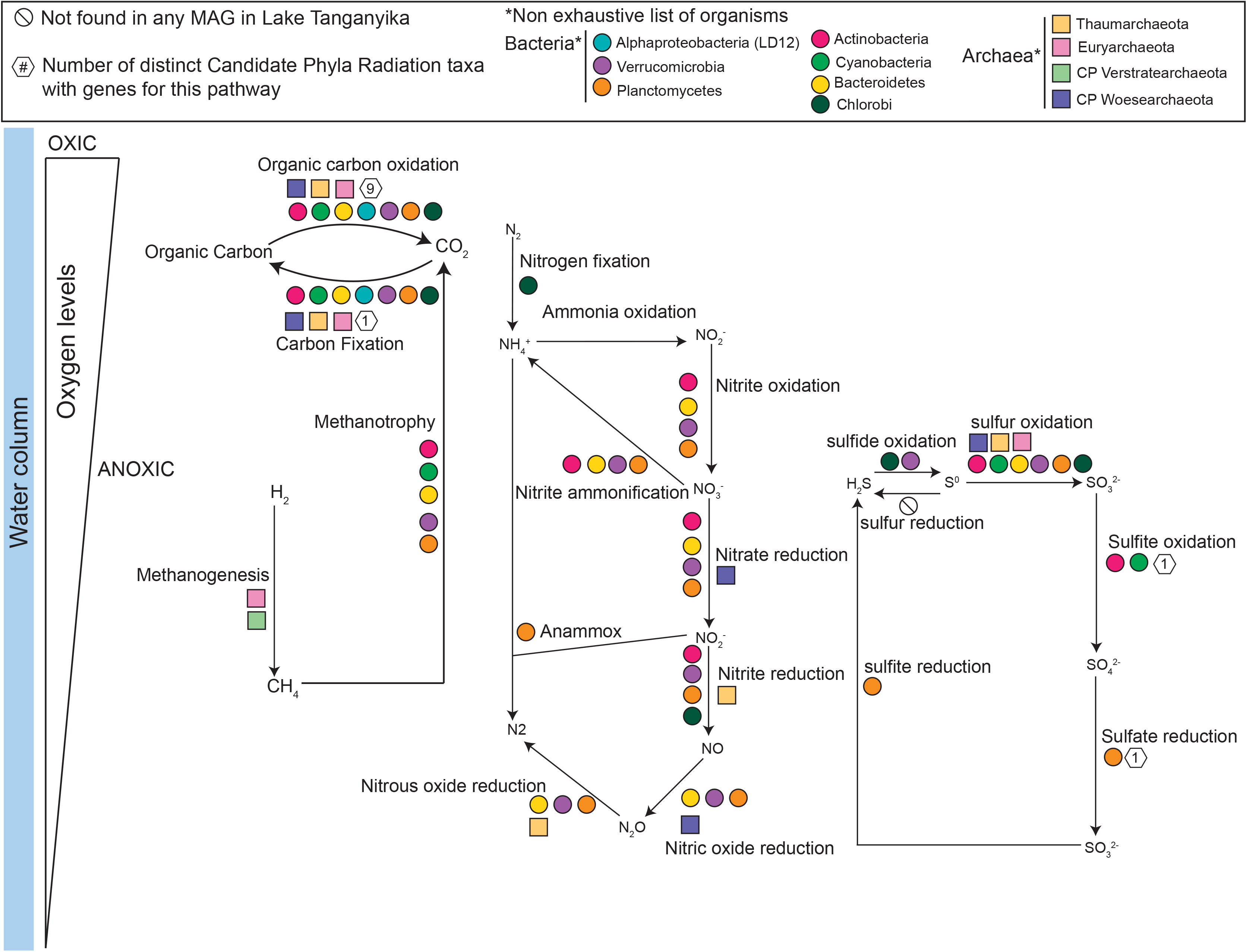
Summary figure showing the role of different microbial taxa in carbon, nitrogen and sulfur cycling in Lake Tanganyika. The list is organisms shown and the reactions are not exhaustive.

To understand the behaviors of carbon, sulfur, and nitrogen biogeochemical cycles in the LT water column, we investigated the depth distribution of different microbially-mediated processes. We observed significant differences amongst the carbon, sulfur, and nitrogen cycles. Processes that were more prominent at sub-oxic and anoxic depths, but still present at low (<5% RAR) from the surface include nitrite oxidation, nitrous oxide reduction, nitrite ammonification, nitrate reduction, methanotrophy, hydrogen oxidation, and hydrogen generation. Processes that were exclusive to the sub-oxic and anoxic depths include methanogenesis, nitrogen fixation, ammonia oxidation, nitrite oxide reduction, nitric oxide reduction, anammox, sulfide oxidation, sulfite reduction, thiosulfate disproportionation into H_2_S and SO_3_^2-^. The potential for use of alternative electron acceptors such as chlorate, metals, arsenate, and selenite were also identified in organisms in sub-oxic and anoxic depths.

Given our earlier finding that the water column was populated with common freshwater taxa in the upper mixed layer, versus less common organisms in the anoxic water column, we wanted to know if the differences were reflected in metabolism of carbon, nitrogen and sulfur across these depths. We identified the metabolic potential of organisms from the lineages Cyanobacteria, Actinobacteria, Alphaproteobacteria (LD12), Verrucomicrobia, Planctomycetes and Bacteroidetes **(Figure 6).** We counted the number of distinct phylogenetic taxa with the potential to conduct a biogeochemical transformation. The potential for nitrogen fixation was not identified in Cyanobacteria. Instead, a small group of microorganisms from 7 distinct taxa which were mostly abundant at sub-oxic and anoxic depths were observed to be capable of nitrogen fixation (Deltaproteobacteria, Kirimatiellacea, Alphaproteobacteria (non LD12), Chloroflexi, Chlorobi, Euryarchaeota, Gammaproteobacteria). The Cyanobacteria in LT all belonged to a Synechoccales, which are known for not being nitrogen fixers[46–49]. The inferred involvement of microbes in biogeochemical cycling across an oxygen gradient such as in LT is summarized in **Figure 7**.

## Discussion

The natural history of LT, along with its environmental characteristics, makes it ideal for examining the role of microbial communities in nutrient and biogeochemical cycling in tropical freshwater lakes. Being an ancient, deep, stratified, and meromictic lake with varying oxygen concentrations, it also serves as a case study for microbial diversity and metabolism in permanently anoxic environments. Here we highlight that microbial diversity and function in LT is distinct between its oxic mixed surface waters, and its anoxic deep waters. We note that although MAGs have key genes to participate in a given process, this does not necessarily mean the corresponding organisms are actively using them all the time. As such, the purpose of our study was to provide an overview of the metabolic potential for microbially-driven biogeochemical processes in LT and to guide future hypotheses for the study of tropical freshwater lakes under stress from a changing climate.

### Comparison of microbiomes between two deep rift valley lakes, Lake Tanganyika and Baikal: Endemism versus Shared lineages

LT (1470m) and Lake Baikal (LB) (1642m) are the world’s two deepest freshwater lakes, yet are drastically different in their water column characteristics. Additionally, both lakes are ancient lakes: LT is approximately 10 million years, and LB is 25 million years. Since few metagenomic studies have been conducted on deep freshwater lakes, a comparison of LT and LB is informative in addressing questions about ancient microbial lineages, evolution, and endemic or shared lineages. As mentioned previously, LT is tropical and has an anoxic bottom layer. LB is located in Siberia, is seasonally ice-covered (twice a year), is dimictic (mixed twice a year), and has an oxygenated bottom layer. Both are ancient and formed geologically through rifts. LB’s microbial community has been studied using metagenomics, from depths ranging from 0-70m during the ice-cover period [6, 50] and from 1250m and 1300m deep collected in March 2018 also during an ice covered period[8]. LT’s microbiome has been studied via 16S rRNA amplicon sequencing in the past [9–11] and in this study via metagenomics. While LT is primarily anoxic below ~100-150m, LB has deep-water currents near the coastal regions which results in oxygenation of deep waters. The regular mixing from surface waters from LB may imply lower endemism, as the “deep water” ecological niche would be more frequently disturbed than in meromictic LT.

On one hand, because both LT and LB are ancient, deep, and rift valley freshwater lakes, we hypothesized that they might have a core microbiome common to both (shared lineages). However, given their contrasting environments (cold vs tropical, oxic vs. anoxic), we expected that environmental characteristics driven by oxygen presence would be major drivers of microbial community structure and function, such as is the case in soils[51], sediments[52], marine oxygen-minimum zones[53]) rather than ecosystem type, especially with the increased importance of nitrogen and sulfur cycling under low-oxygen conditions. In our study for example, we find taxa such as sulfur cycling Desulfobacterota only found in LT and not in Baikal, likely reflecting that these H2S producers are found in low oxygen environments.

### Metabolic connections across vertical gradients in LT

Our metabolic function analyses show the distribution of microbially-driven carbon, nitrogen, and sulfur cycling across a depth gradient in LT. The carbon cycle of LT has mostly been investigated from the perspective of primary production, and methane cycling. It has been suggested that anaerobic methane oxidation contributes to decreasing the amount of methane in the anoxic zone of LT [2]. Methane is a greenhouse gas that is important in LT, particularly with respect to stratification. Because methane is more abundant below the oxycline, a shift in the oxycline could lead to methane release to the atmosphere via ebullition. Bacterial activity is thought to be a source of methane in nearby (~ 470 km away) Lake Kivu [54]. Major findings in methanogenesis in aquatic ecosystems now bring light to previously unrecognized biological sources and sinks of methane [55, 56]. We and others [2] have found methanogenic Archaea in LT.

Nitrogen cycling has been well studied in LT, mostly focused on nitrogen inputs and outputs, and from a chemical and biogeochemical perspective. For example, atmospheric deposition and nitrogen from river inflow contribute much of the nitrogen to LT[57]. Additionally, it is shown that incoming nitrogen in the form of nitrate is quickly used by organisms in the surface waters, and the upper water column is often nitrate depleted. Profiles of nitrogen (NH_4_^+^, NO_2_^-^, NO_3_^-^) in LT show that NH3 accumulated from 200m to 1200m, NO_2_^-^ is usually overall low throughout the water column, and NO_3_^-^ is low at the surface, peaks at the oxycline, and declines where NH_4_^+^ increases[40]. Another way that NH_4_^+^ can be supplemented to the water column is through nitrogen fixation. In freshwater systems, it is typically thought that this process is conducted by Cyanobacteria, particularly the heterocystous *Anabaena flos-aquae*[58]. In Lake Mendota, a eutrophic temperate freshwater lake, metagenomic data has shown that a third of nitrogen fixation genes originate from Cyanobacteria, and the remaining from Betaproteobacteria and Gammaproteobacteria[44]. However, we were surprised to find that nitrogen fixation was not identified in any LT Cyanobacteria. Benthic nitrogen fixers (cyanobacteria and diatoms with endosymbionts) are dominant in the littoral zone of LT and can provide as much as 30% of N in nearby lake Malawi[59]. Rather, from the metagenomic data of free-living planktonic bacteria, nitrogen fixation was identified to potentially be perfomed in other groups such as Deltaproteobacteria, Kirimatiellacea, Alphaproteobacteria, Chloroflexi, Chlorobi, Euryarchaeota, and Gammaproteobacteria. Similar results have been observed in Trout Bog Lake, a stratified humic lake where the nitrogen fixers were more diverse and also included Chlorobi [44], like in LT. From the chemical evidence, we hypothesized that the oxycline is a hotspot for nitrite oxidation, and denitrification occurs below the oxycline. Nitrite oxidation pathways in LT MAGs show that many organisms are potentially able to perform the reaction, however the *nxr* enzyme may act reversibly (NO_2_^-^ to NO_3_^-^ and vice-versa). Denitrification processes (nitrate reduction, nitrite reduction, nitric oxide reduction, and nitrous oxide reduction) have similar representation across all 3 depths. A study of nitrogen cycling processes in LT[60] identified very low dissolved inorganic nitrogen concentrations in the euphotic zone, and proposed that very active N cycling must be occurring in the water column to prevent this accumulation. Another process leading to low dissolved inorganic nitrogen concentration is anammox. Incubations with ^15^N labelled nitrate have showed that anammox was active in the 100-110m water depth, and was comparable to rates observed in marine oxygen minimum zones [11]. Our detection of Planctomycetes bacteria capable of anammox (Brocadiales) supports this conclusion.

Sulfur cycling in freshwater lakes has received less attention compared to carbon, nitrogen, and phosphorus. Profiles from LT show a clear increase in H2S below the oxycline, stabilizing at a concentration of ~30μM H2S from 300m to 1200m depth [40]. Biologically, H2S is produced via sulfite reduction, thiosulfate disproportionation into H_2_S and SO_3_^2-^, sulfur reduction, all of which were identified exclusively in the sub-oxic and anoxic depths. Given the use of sulfur based electron acceptors as alternatives to oxygen in anoxic systems, sulfur cycling has been documented to increase in importance with lower-oxygen availability. As one of the critical compounds for life, sulfur availability and processing is essential to support cellular functions. In LT, where a large portion of the water column is anoxic, sulfur cycling is expected to support primary productivity in the upper water column, and have an overall important effect on lake ecology and biogeochemistry[61]. While the abundance of sulfur cycling organisms suggests an active sulfur cycle in LT, intriguingly, the abundance of organisms mediating the oxidation of reduced sulfur compounds (H_2_S, S^0^, S_2_O_4_^2-^) far exceeded the abundance of organisms mediating the reduction of sulfur compounds (SO_3_^2-^, SO_4_^2-^, S^0^). While this discrepancy may arise from increased enzymatic activity of sulfate/sulfite and sulfur reducing organisms, it may also be associated with the geochemistry of the deep waters in LT. Being a rift valley lake, it is home to hydrothermal vents, in which reduced compounds and gases from the Earth’s crust including H_2_, H_2_S, and CH_4_ may be produced [62, 63]. Around these hydrothermal vents, white and brown microbials mats have previously been identified[63] and microbial life is abundant[64]. In marine environments, *Beggiatoa* mats are associated with sulfide oxidation[65], and Zetaproteobacteria are often the primary iron oxidizers in iron-rich systems, creating brown mats[66].

## Conclusions

Our study provides genomic evidence for the capabilities of Bacteria and Archaea in tropical freshwater biogeochemical cycling and describes links between spatial distribution of organisms and biogeochemical processes. These processes are known to impact critical food webs that are renowned for their high biodiversity and that serve as important protein sources for local human populations, as the shoreline of LT is shared by four African countries. As a permanently stratified lake, LT also offers a window into the ecology and evolution of microorganisms in tropical freshwater environments. Our study encompassed the lake’s continuous vertical redox gradient, and microbial communities were dominated by core freshwater taxa at the surface and by Archaea and uncultured Candidate Phyla in the deeper anoxic waters. The high prominence of Archaea making up to 30% of the community composition abundance at lower depths further highlighted this contrast. While freshwater lakes are often limited by phosphorus [67], tropical freshwater lake are frequently nitrogen-limited[68]. Our microbe-centric analyses reveal that abundant microbes in LT play important roles in nitrogen transformations thay may remove fixed nitrogen from the water column, or fix nitrogen that may be upwelled and replenish the productive surface waters in bioavailable forms of fixed nitrogen.

Tropical lakes are abundant on earth, yet understudied both in the limnological literature and in the context of microbial metabolism and chemistry. Yet, it is critically important to understand the biogeochemistry of tropical lakes, as lake ecosystem health is critical to the livelihood of millions of people, for example in LT[69, 70]. With increasing global temperatures, extremely deep lakes such as LT will likely experience increased stratification, lower mixing, and increased anoxia [71]. We provide the most comprehensive metagenomics-based genomic study to date of microbial community metabolism in an anoxic tropical freshwater lake which is an essential foundation for future work on environmental adaptation, food webs, and nutrient cycling. Overall, our study will enable the continued integration of aquatic ecology, ‘omics’ data, and biogeochemistry in freshwater lakes, that is key to a holistic understanding of ongoing global change and its impact on surface freshwater resources.

## Acknowledgements

We thank the University of Wisconsin – Office of the Vice Chancellor for Research and Graduate Education, University of Wisconsin – Department of Bacteriology, and University of Wisconsin – College of Agriculture and Life Sciences for their support. The Lake Tanganyika project was supported by the United States National Science Foundation (DEB-1030242 to P.B.M and DEB-0842253 to Y.V.). We are thankful to the Tanzania Commission for Science and Technology (COSTECH) for providing the research permits to collect the samples. This research was also supported by the U.S. Department of Energy Joint Genome Institute (JGI) through a JGI-Community Science Program Award to K.D.M (Proposal ID: CSP 2796). The work conducted by the U.S. Department of Energy Joint Genome Institute, a DOE Office of Science User Facility, is supported by the Office of Science of the U.S. Department of Energy under Contract No. DE-AC02-05CH11231. K.D.M. received funding from the United States National Science Foundation via an INSPIRE award (DEB-1344254) and the Wisconsin Alumni Research Foundation at UW-Madison. B.M.K was supported by funding from the Leibniz Institute for Freshwater Ecology and Inland Fisheries’ International Postdoctoral Research Fellowship and from the German Research Foundation through the LimnoScenES project (AD 91/22-1). S.C.B. was supported by a NSF-REU award for Summer 2020. P.Q.T is supported by the Natural Sciences and Engineering Research Council of Canada (NSERC). I (P.Q.T) am thankful to our colleagues of Anantharaman labs and the McMahon labs for valuable feedback throughout the project including A. Schmidt for teaching me geographic spatial analysis, and Z. Zhou for co-developing our metagenome analysis pipeline in the Anantharaman lab. We thank all anonymous reviewers for their comments.

## Data availability

The MAGs can be accessed on NCBI BioProject ID PRJNA523022. The genomes will be officially released on NCBI Genbank upon publication. All MAGs are also publicly available on the Open Science Framework: https://osf.io/pmhae/. The 24 raw and assembled metagenomes are available on the Integrated Microbial Genomes & Microbiomes (IMG/M) portal using the following IMG Genome ID’s: 3300020220, 3300020083, 3300020183, 3300020200, 3300021376, 3300021093, 3300021091, 3300020109, 3300020074, 3300021092, 3300021424, 3300020179, 3300020193, 3300020204, 3300020221, 3300020196, 3300020190, 3300020197, 3300020222, 3300020214, 3300020084, 3300020198, 3300020603, 3300020578. An interactive version of the Archaeal and Bacteria trees can be accessed at iTOL at: https://itol.embl.de/shared/patriciatran. Code to generate the figures, and access to the custom HMM for the 16 ribosomal proteins and the metabolic genes is available at https://github.com/patriciatran/LakeTanganyika/.

## Supplementary Figures

**Supplementary Figure 1.** Overview of all samples from this study. Chlorophyll *a*, conductivity, dissolved Oxygen (DO), and temperature was collected from 2010 to 2013. Secchi depth data are available from 2012 and 2015. Metagenome samples were collected in 2015.

**Supplementary Figure 2.** Environmental Data Profiles from 2010-2013. Dots represent the thermocline depth at each sampling date.

**Supplementary Figure 3.** A. Secchi depths in Lake Tanganyika. B. Kd coefficient in Lake Tanganyika, calculated by 1.7 divided by the Secchi depth in meters. Location Lat −4.89, Long 29.59 is approximately where the metagenome: KigOffshore Cast was taken, which consists of samples KigOff0, KigOff40, KigOff80 and KigOff120. Based on this trend of the Secchi depths presented in this figure, we estimate that all three metagenome KigOff40, 80, and 120 would be below the Secchi Depth.

**Supplementary Figure 4.** Soluble reactive phosphorus (SRP) concentrations, which is a limnological indicator of phosphorus (blue), and nitrate concentration (red) in Lake Tanganyika. Data was collected on July 15, 2015 in Lake Tanganyika at the Kigoma and Mahale station. The peak in nitrate occurred between 50 and 65m.

**Supplementary Figure 5.** Chlorophyll *a* increases with up (down to 150m depth sampling) peak generally around 120m. Over one year, chlorophyll *a* increases across all depth (points shift towards the right).

**Supplementary Figure 6. A.** Comparison of manual RP16 taxonomy versus the GTDB-tk automated taxonomy, and distribution of completeness levels on for each taxonomy group. **B.** Distribution of completeness values among categories. The colors in panel B is the legend for the colors in the bar plot of panel A. There were matches across the range of completeness values (“Match”). However, when there is a mismatch, it is not always because the genome is less complete.

**Supplementary Figure 7. A.** Rank abundance curve overall for all samples **B.** Rank Abundance curve by station **C.** Rank abundance curve for all samples, grouped by sample depth (meters).

**Supplementary Figure 8.** Comparison of the taxonomic diversity between Lake Baikal and Lake Tanganyika. **A.** Number of MAGS shared between the two lakes and unique to either lake. **B.** Comparison of taxonomic diversity at the phylum-level between the two lakes.

**Supplementary Figure 9.** Heatmap showing the genes involved in carbon cycling found in the MAGs.

**Supplementary Figure 10.** Heatmap showing the genes involved in nitrogen cycling in the MAGs.

**Supplementary Figure 11**. Heatmap showing the genes involved in sulfur cycling found in the MAGs.

**Supplementary Figure 12**. Heatmap showing the genes involved in metal biogeochemical cycling found in the MAGs, for other types of metabolism.

**Supplementary Figure 13.** Distribution of carbohydrate active enzymes in LT. Bar plot showing the Cazyme densities (number of cazyme hits divided by genome size in bp, in percentage), and number of hits to each CAZyme category: AA, CBM (carbohydrate binding module), CE, cohesin, GH (glycoside hydrolases), GT (glycosyl transferase), PL (polysaccharide lyase). The taxonomic groups are on the y-axis, and the number of MAGs corresponding to each group is listed in parentheses. Vertical grey bars highlight grouping of Verrucomicrobia (which are split into Orders), Alphaproteobacteria, and Actinobacteria.

**Supplementary Figure 14.** Single-gene phylogeny of amoA to identify (MAFFT, RAxML) different groups of nitrite-oxidizing bacteria or nitrate-reducing bacteria.

**Supplementary Figure 15.** Single-gene phylogeny of nxrA to identify (MAFFT, RAxML) different groups of nitrite-oxidizing bacteria or nitrate-reducing bacteria.

**Supplementary Figure 16.** Expanded figure showing the comammox Nitrospira from Lake Tanganyika in the context of references comammox genomes. **A.** Concatenated RP16 gene phylogeny of Nitrospira genomes from LT (renamed LTC-1 and LTC-2) and other environments (Accession Numbers in parentheses). **B.** Presence (filled boxes) and absence (empty boxes) of selected genes involved in ammonia and nitrite oxidation. The presence of comammox is based on the presence of both amo and nxr genes. ammonium transporters and urea utilization.

## Supplementary Tables

**Table S1** Description of the 24 metagenome samples collected in LT

**Table S2** Genome characteristics for the MAGs recovered in LT.

**Table S3** Whole-genome pairwise ANI values between the 3 Candidatus Tanganyikabacteria MAGs against Cyanobacteria, MAGs from freshwater lakes, Melainabacteria, Margulisbacteria, Blackallbacteria references for a total of 309 genomes.

**Table S4** Relative Abundance of Reads mapped (RAR) of MAGs across the 24 metagenomic samples.

**Table S5** Genome-level ANI comparisons of taxonomic groups present in the deepest samples from LT and LB.

**Table S6** Results summarizing the METABOLIC output, showing the number of distinct MAGs involved in each reaction, and their taxonomic identities.

**Table S7** Presence and absence of genes based on the hits from 143 HMMs searches.

**Table S8** Identification of carbohydrate active enzymes (CAZYmes) in MAGs from LT.

## Supplementary Text

Section 1. Detailed Methods for producing the sampling map.

Section 2. Detailed methods to curate the nxrA gene using a single-gene phylogeny, to distinguish between nitrite oxidizers and nitrate reducers

Section 3. Detailed Methods to identify Archeal amoA gene, using a singe-gene phylogeny of Bacterial and Archeal amoA gene, and using BLAST from known Archeal amoABC gene against the Lake Tanganyika Archaeal MAGs.

Section 4. Detailed Methods to generate the Nitrospira Tree (RP16).

## Supplementary Material

**Supplementary Material 1** Newick File for concatenated ribosomal protein phylogeny Archaeal Tree (RAXML)

**Supplementary Material 2** Newick File for concatenated ribosomal protein phylogeny Bacterial Tree (RAXML)

**Supplementary Material 3** GTDB-tk Generated concatenated gene phylogeny using FastTree (Archaea)

**Supplementary Material 4** GTDB-tk generated concatenated gene phylogeny using FastTree, and 120 marker genes (Bacteria).

## References

1. Alin SR, Johnson TC. Carbon cycling in large lakes of the world: A synthesis of production, burial, and lake-atmosphere exchange estimates: CARBON CYCLING IN LARGE LAKES. Global Biogeochemical Cycles 2007; 21: n/a-n/a.

2. Durisch-Kaiser E, Schmid M, Peeters F, Kipfer R, Dinkel C, Diem T, et al. What prevents outgassing of methane to the atmosphere in Lake Tanganyika? Journal of Geophysical Research 2011; 116.

3. Takahashi T, Koblmüller S. The Adaptive Radiation of Cichlid Fish in Lake Tanganyika: A Morphological Perspective. International Journal of Evolutionary Biology 2011; 2011: 1–14.

4. Salzburger W. Understanding explosive diversification through cichlid fish genomics. Nat Rev Genet 2018; 19: 705–717.

5. Corman JR, McIntyre PB, Kuboja B, Mbemba W, Fink D, Wheeler CW, et al. Upwelling couples chemical and biological dynamics across the littoral and pelagic zones of Lake Tanganyika, East Africa. Limnology and Oceanography 2010; 55: 214–224.

6. Cabello-Yeves PJ, Zemskaya TI, Rosselli R, Coutinho FH, Zakharenko AS, Blinov VV, et al. Genomes of novel microbial lineages assembled from the sub-ice waters of Lake Baikal. Applied and Environmental Microbiology 2017; AEM.02132–17.

7. Cabello-Yeves PJ, Zemskaya TI, Rosselli R, Coutinho FH, Zakharenko AS, Blinov VV, et al. Genomes of Novel Microbial Lineages Assembled from the Sub-Ice Waters of Lake Baikal. Appl Environ Microbiol 2018; 84.

8. Cabello-Yeves PJ, Zemskaya TI, Zakharenko AS, Sakirko MV, Ivanov VG, Ghai R, et al. Microbiome of the deep Lake Baikal, a unique oxic bathypelagic habitat. Limnology and Oceanography 2019.

9. De Wever A. Spatio-temporal dynamics in the microbial food web in Lake Tanganyika. University of Gent 2006; 1–169.

10. Pirlot S, Unrein F, Descy J-P, Servais P. Fate of heterotrophic bacteria in Lake Tanganyika (East Africa): Fate of bacteria in Lake Tanganyika. FEMS Microbiology Ecology 2007; 62: 354–364.

11. Schubert CJ, Durisch-Kaiser E, Wehrli B, Thamdrup B, Lam P, Kuypers MMM. Anaerobic ammonium oxidation in a tropical freshwater system (Lake Tanganyika). Environmental Microbiology 2006; 8: 1857–1863.

12. Shade A, Kent AD, Jones SE, Newton RJ, Triplett EW, McMahon KD. Interannual dynamics and phenology of bacterial communities in a eutrophic lake. Limnology and Oceanography 2007; 52: 487–494.

13. Nurk S, Meleshko D, Korobeynikov A, Pevzner PA. metaSPAdes: a new versatile metagenomic assembler. Genome Res 2017; 27: 824–834.

14. Kang DD, Froula J, Egan R, Wang Z. MetaBAT, an efficient tool for accurately reconstructing single genomes from complex microbial communities. PeerJ 2015; 3: e1165.

15. Kang DD, Li F, Kirton E, Thomas A, Egan R, An H, et al. MetaBAT 2: an adaptive binning algorithm for robust and efficient genome reconstruction from metagenome assemblies. PeerJ 2019; 7: e7359.

16. Wu Y-W, Simmons BA, Singer SW. MaxBin 2.0: an automated binning algorithm to recover genomes from multiple metagenomic datasets. Bioinformatics 2016; 32: 605–607.

17. Sieber CMK, Probst AJ, Sharrar A, Thomas BC, Hess M, Tringe SG, et al. Recovery of genomes from metagenomes via a dereplication, aggregation and scoring strategy. Nature Microbiology 2018; 3: 836–843.

18. Olm MR, Brown CT, Brooks B, Banfield JF. dRep: a tool for fast and accurate genomic comparisons that enables improved genome recovery from metagenomes through dereplication. The ISME Journal 2017; 1–5.

19. Parks DH, Imelfort M, Skennerton CT, Hugenholtz P, Tyson GW. CheckM: assessing the quality of microbial genomes recovered from isolates, single cells, and metagenomes. Genome research 2015; 25: 1043–55.

20. The Genome Standards Consortium, Bowers RM, Kyrpides NC, Stepanauskas R, Harmon-Smith M, Doud D, et al. Minimum information about a single amplified genome (MISAG) and a metagenome-assembled genome (MIMAG) of bacteria and archaea. Nature Biotechnology 2017; 35: 725–731.

21. Bushnell B. BBMAP. https://jgi.doe.gov/data-and-tools/bbtools/bb-tools-user-guide/bbmap-guide/..

22. Hyatt D, Chen G-L, LoCascio PF, Land ML, Larimer FW, Hauser LJ. Prodigal: prokaryotic gene recognition and translation initiation site identification. BMC Bioinformatics 2010; 11.

23. Anantharaman K, Brown CT, Hug LA, Sharon I, Castelle CJ, Probst AJ, et al. Thousands of microbial genomes shed light on interconnected biogeochemical processes in an aquifer system. Nature Communications 2016; 7: 13219.

24. Eddy SR. Accelerated profile HMM searches. PLoS Computational Biology 2011; 7.

25. Hug LA, Baker BJ, Anantharaman K, Brown CT, Probst AJ, Castelle CJ, et al. A new view of the tree of life. Nature Microbiology 2016; 1: 1–6.

26. Brown AMV, Howe DK, Wasala SK, Peetz AB, Zasada IA, Denver DR. Comparative genomics of a plant-parasitic nematode endosymbiont suggest a role in nutritional symbiosis. Genome Biology and Evolution 2015; 7: 2727–2746.

27. Katoh K, Standley DM. MAFFT multiple sequence alignment software version 7: Improvements in performance and usability. Molecular Biology and Evolution 2013; 30: 772–780.

28. Miller MA, Pfeiffer W, Schwartz Terri. Creating the CIPRES Science Gateway for Inference of Large Phylogenetic Trees. Proceedings of the Gateway Computing Environments Workshop. 2010. New Orleans, LA, pp 1–8.

29. Parks DH, Chuvochina M, Waite DW, Rinke C, Skarshewski A, Chaumeil P-A, et al. A standardized bacterial taxonomy based on genome phylogeny substantially revises the tree of life. Nature Biotechnology 2018; 36: 996–1004.

30. Newton RJ, Jones SE, Eiler A, McMahon KD, Bertilsson S. A guide to the natural history of freshwater lake bacteria. Microbiology and molecular biology reviews: MMBR. 2011.

31. Rohwer RR, Hamilton JJ, Newton RJ, McMahon KD. TaxAss: Leveraging a Custom Freshwater Database Achieves Fine-Scale Taxonomic Resolution. mSphere 2018; 3.

32. Soo RM, Hemp J, Parks DH, Fischer WW, Hugenholtz P. On the origins of oxygenic photosynthesis and aerobic respiration in Cyanobacteria. Science 2017; 355: 1436–1440.

33. Linz AM, He S, Stevens SLR, Anantharaman K, Robin R. Connections between freshwater carbon and nutrient cycles revealed through. 2018; 221.

34. Bendall ML, Stevens SL, Chan L-K, Malfatti S, Schwientek P, Tremblay J, et al. Genomewide selective sweeps and gene-specific sweeps in natural bacterial populations. The ISME Journal 2016; 10: 1589–1601.

35. Jain C, Rodriguez-R LM, Phillippy AM, Konstantinidis KT, Aluru S. High throughput ANI analysis of 90K prokaryotic genomes reveals clear species boundaries. Nature Communications 2018; 9.

36. Soo RM, Skennerton CT, Sekiguchi Y, Imelfort M, Paech SJ, Dennis PG, et al. An Expanded Genomic Representation of the Phylum Cyanobacteria. Genome Biology and Evolution 2014; 6: 1031–1045.

37. Zhou Z, Tran P, Liu Y, Kieft K, Anantharaman K. METABOLIC: A scalable high-throughput metabolic and biogeochemical functional trait profiler based on microbial genomes. bioRxiv 2019; 761643.

38. Zhang H, Yohe T, Huang L, Entwistle S, Wu P, Yang Z, et al. dbCAN2: a meta server for automated carbohydrate-active enzyme annotation. Nucleic Acids Research 2018; 46: W95–W101.

39. Mukherjee S, Stamatis D, Bertsch J, Ovchinnikova G, Katta HY, Mojica A, et al. Genomes OnLine database (GOLD) v.7: updates and new features. Nucleic Acids Res 2019; 47: D649–D659.

40. Edmond JM, Stallard RF, Craig H, Craig V, Weiss RF, Coulter GW. Nutrient chemistry of the water column of Lake Tanganyika. Limnology and Oceanography 1993; 38: 725–738.

41. Verburga P, Hecky RE. The physics of the warming of Lake Tanganyika by climate change. Limnology and Oceanography 2009; 54: 2418–2430.

42. Järvinen M, Salonen K, Sarvala J, Vuorio K, Virtanen A. The stoichiometry of particulate nutrients in Lake Tanganyika – implications for nutrient limitation of phytoplankton. Hydrobiologia 1999; 407: 81–88.

43. Ehrenfels B, Bartosiewicz M, Mbonde AS, Baumann KBL, Dinkel C, Junker J, et al. Thermocline depth and euphotic zone thickness regulate the abundance of diazotrophic cyanobacteria in Lake Tanganyika. 2020. Biogeochemistry: Limnology.

44. Linz AM, He S, Stevens SLR, Anantharaman K, Rohwer RR, Malmstrom RR, et al. Freshwater carbon and nutrient cycles revealed through reconstructed population genomes. PeerJ 2018; 6:e6075.

45. Martinez-Garcia M, Brazel DM, Swan BK, Arnosti C, Chain PSG, Reitenga KG, et al. Capturing single cell genomes of active polysaccharide degraders: An unexpected contribution of verrucomicrobia. PLoS ONE 2012; 7: 1–11.

46. Damrow R, Maldener I, Zilliges Y. The Multiple Functions of Common Microbial Carbon Polymers, Glycogen and PHB, during Stress Responses in the Non-Diazotrophic Cyanobacterium Synechocystis sp. PCC 6803. Front Microbiol 2016; 7.

47. Paerl HW, Otten TG. Duelling ‘CyanoHABs’: unravelling the environmental drivers controlling dominance and succession among diazotrophic and non-N2-fixing harmful cyanobacteria. Environmental Microbiology 2016; 18: 316–324.

48. Raymond J, Siefert JL, Staples CR, Blankenship RE. The Natural History of Nitrogen Fixation. Mol Biol Evol 2004; 21: 541–554.

49. Berman-Frank I, Lundgren P, Falkowski P. Nitrogen fixation and photosynthetic oxygen evolution in cyanobacteria. Research in Microbiology 2003; 154: 157–164.

50. Cabello-Yeves PJ, Ghai R, Mehrshad M, Picazo A, Camacho A, Rodriguez-valera F. Reconstruction of diverse verrucomicrobial genomes from metagenome datasets of freshwater reservoirs. Frontiers in Microbiology 2017; 8.

51. Hansel CM, Fendorf S, Jardine PM, Francis CA. Changes in Bacterial and Archaeal Community Structure and Functional Diversity along a Geochemically Variable Soil Profile. Appl Environ Microbiol 2008; 74: 1620–1633.

52. Edlund A, Hårdeman F, Jansson JK, Sjöling S. Active bacterial community structure along vertical redox gradients in Baltic Sea sediment. Environmental Microbiology 2008; 10: 2051–2063.

53. Beman JM, Carolan MT. Deoxygenation alters bacterial diversity and community composition in the ocean’s largest oxygen minimum zone. Nature Communications 2013; 4: 2705.

54. Schoell M, Tietze K, Schoberth SM. Origin of methane in Lake Kivu (East-Central Africa). Chemical Geology 1988; 71: 257–265.

55. Bogard MJ, del Giorgio PA, Boutet L, Chaves MCG, Prairie YT, Merante A, et al. Oxic water column methanogenesis as a major component of aquatic CH4 fluxes. Nature Communications 2014; 5.

56. Vanwonterghem I, Evans PN, Parks DH, Jensen PD, Woodcroft BJ, Hugenholtz P, et al. Methylotrophic methanogenesis discovered in the archaeal phylum Verstraetearchaeota. Nature Microbiology 2016; 1.

57. Gao Q, Chen S, Kimirei IA, Zhang L, Mgana H, Mziray P, et al. Wet deposition of atmospheric nitrogen contributes to nitrogen loading in the surface waters of Lake Tanganyika, East Africa: a case study of the Kigoma region. Environmental Science and Pollution Research 2018; 25: 11646–11660.

58. Chale FMM. Inorganic Nutrient Concentrations and Chlorophyll in the Euphotic Zone of Lake Tanganyika. Hydrobiologia 2004; 523: 189–197.

59. Higgins SN, Hecky RE, Taylor WD. Epilithic nitrogen fixation in the rocky littoral zones of Lake Malawi, Africa. Limnology and Oceanography 2001; 46: 976–982.

60. Brion N, Nzeyimana E, Goeyens L, Nahimana D, Tungaraza C, Baeyens W. Inorganic Nitrogen Uptake and River Inputs in Northern Lake Tanganyika. Journal of Great Lakes Research 2006; 32: 553–564.

61. Norici A, Hell R, Giordano M. Sulfur and primary production in aquatic environments: an ecological perspective. Photosynth Res 2005; 86: 409–417.

62. Botz RW, Stoffers P. Light hydrocarbon gases in Lake Tanganyika hydrothermal fluids (East-Central Africa). Chemical Geology 1993; 104: 217–224.

63. Tiercelin J-J, Pflumio C, Castrec M, Boulégue J, Gente P, Rolet J, et al. Hydrothermal vents in Lake Tanganyika, East African, Rift system. Geology 1993; 21: 499–502.

64. Elsgaard L, Prieur D. Hydrothermal vents in Lake Tanganyika harbor spore-forming thermophiles with extremely rapid growth. Journal of Great Lakes Research 2011; 37: 203–206.

65. Preisler A, de Beer D, Lichtschlag A, Lavik G, Boetius A, Jørgensen BB. Biological and chemical sulfide oxidation in a Beggiatoa inhabited marine sediment. The ISME Journal 2007; 1: 341–353.

66. McAllister SM, Moore RM, Gartman A, Luther GW, Emerson D, Chan CS. The Fe(II)-oxidizing Zetaproteobacteria: historical, ecological and genomic perspectives. FEMS Microbiol Ecol 2019; 95.

67. Carpenter SR. Phosphorus control is critical to mitigating eutrophication. Proceedings of the National Academy of Sciences 2008; 105: 11039–11040.

68. Jr WML. Causes for the high frequency of nitrogen limitation in tropical lakes. SIL Proceedings, 1922-2010 2002; 28: 210–213.

69. De Keyzer ELR, Masilya Mulungula P, Alunga Lufungula G, Amisi Manala C, Andema Muniali A, Bashengezi Cibuhira P, et al. Local perceptions on the state of the pelagic fisheries and fisheries management in Uvira, Lake Tanganyika, DR Congo. Journal of Great Lakes Research 2020.

70. MÖLSÄ, Hannu. Management of Fisheries on Lake Tanganyika Challenges for Research and the Community. 2008. University of Kuopio.

71. Foley B, Jones ID, Maberly SC, Rippey B. Long-term changes in oxygen depletion in a small temperate lake: effects of climate change and eutrophication. Freshwater Biology 2012; 57: 278–289.

