## Supplementary Text for "Depth-discrete metagenomics reveals the roles of microbes in biogeochemical cycling in the tropical freshwater Lake Tanganyika"

### Detailed Methods for producing the sampling map

The sample map (**Figure 1**) was created using ArcMap version 9 (January 2020). The basemaps were accessed from Natural Earth (https://www.naturalearthdata.com/) in January 2020. Large scale data (1:10m) for countries, lakes, rivers, major cities, and graticules at 1degree intervals were downloaded. The shapefile for the 24 samples was loaded. For the main map, we chose a region-specific projection, the Africa Albers Equal Area Conic. For the inset small-scale data (1:110m) countries data was downloaded and projected unto a vertical spherical projection.

### Detailed methods to curate the nxrA gene using a single-gene phylogeny, to distinguish between nitrite oxidizers and nitrate reducers.

Nitrite oxidoreductase alpha subunit (nxrA) can be tricky to identify its function from gene calling only, because it is involved in both nitrite oxidation to nitrate, and nitrate reduction to nitrite. As such, one method to parse the function of nxrA is to create a single-gene nxrA tree, which will be split into 2 groups of known nitrite oxidizers, and known nitrate reducers. To do this, we downloaded 25 reference sequences from NCBI RefSeq with predicted nxrA sequences from LT (based on metabolic HMMs). We used MAFFT for sequence alignment, and RAxML-HPC on the CIPRES server to run analyses.

### amoA gene: Differentiating between pmoA and amoA, and detection of Archaeal amoABC subunits in Archeal MAGs

We annotated pmoA using custom HMMs as described above. amoA and pmoA can be difficult to distinguish using gene calling methods. Therefore, we selected sequences annotated as "pmoA" (amoA-like sequences) and built a single-gene phylogeny using references from Alves et al. 2018[1]. We first called the protein coding sequences using Prodigal[2] on the File 1 named "AamoA.db_an96.aln_tax.annotated.fasta". Then we aligned the sequences using MAFFT[3] in Geneious Prime, and applied 90% gap masking. We used RAxML[4] to build a comprehensive tree. Based on the phylogenetic position of the sequences, we delineated amoA and pmoA sequences. We noted that the pmoA custom HMM did not pick up any Archaeal sequences.

To identify archaeal ammonia oxidizers in LT, we searched the amoA, amoB and amoC subunits of Candidatus Nitrosophaera gargensis Ga9.2 (Phylum Thaumarchaeota, Kingdom Crenarchaeota, Class Nitrososphaeria) using BLASTp 2.2.31[5]. We identified 5 Thaumarchaeota MAG with either subunits a, b or c, and 3 Thaumarchaeota MAG with all subunits (amoABC).

### Detailed Methods to Generate the Nitrospira Tree

We downloaded genomes from the Nitrospira taxonomy on NCBI RefSeq, and created a concatenated gene phylogeny using RP16. Nine Rokubacteria genomes was used as the outgroup. We used MAFFT[3] and RAxML[4] on the CIPRES[6] server to generate the tree. The two Nitrospira MAGs from Lake Tanganyika fell into class IIA.

**Cited Literature:**


1. Alves RJE, Minh BQ, Urich T, von Haeseler A, Schleper C. Unifying the global phylogeny and environmental distribution of ammonia-oxidising archaea based on amoA genes. *Nature Communications* 2018; **9**: 1517.
2. Hyatt D, Chen G-L, LoCascio PF, Land ML, Larimer FW, Hauser LJ. Prodigal: prokaryotic gene recognition and translation initiation site identification. *BMC Bioinformatics* 2010; **11**.
3. Katoh K, Standley DM. MAFFT multiple sequence alignment software version 7: Improvements in performance and usability. *Molecular Biology and Evolution* 2013; **30**: 772–780.
4. Liu K, Linder CR, Warnow T. RAxML and FastTree: Comparing Two Methods for Large-Scale Maximum Likelihood Phylogeny Estimation. *PLoS ONE* 2011; **6**: e27731.
5. Altschul SF, Gish W, Miller W, Myers EW, Lipman DJ. Basic local alignment search tool. *J Mol Biol* 1990; **215**: 403–410.
6. Miller MA, Pfeiffer W, Schwartz Terri. Creating the CIPRES Science Gateway for Inference of Large Phylogenetic Trees. *Proceedings of the Gateway Computing Environments Workshop*. 2010. New Orleans, LA, pp 1–8.
