## Supplementary material for "Depth-discrete metagenomics reveals the roles of microbes in biogeochemical cycling in the tropical freshwater Lake Tanganyika": Figure S1

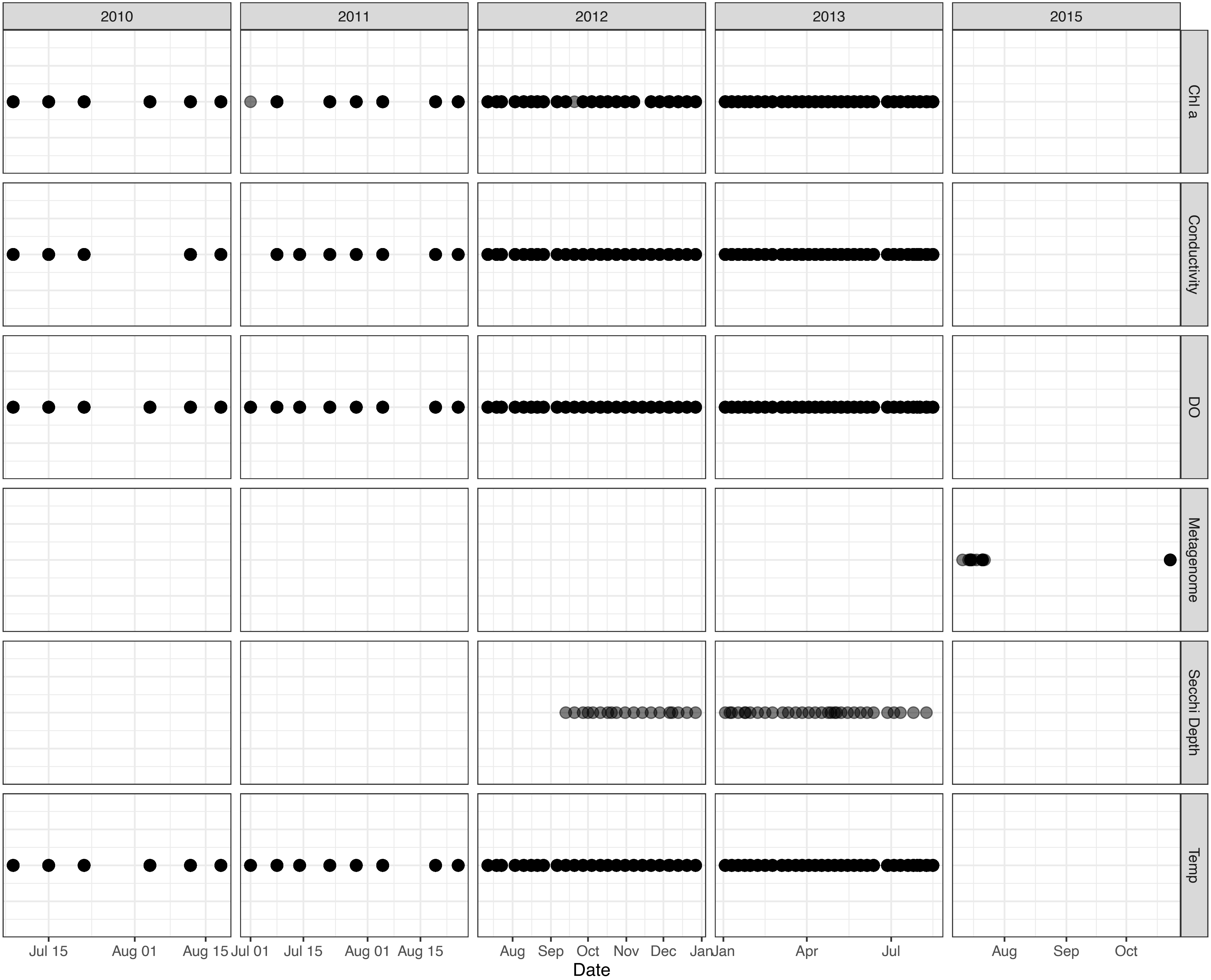


**Supplementary Figure 1.** Overview of all samples from this study. Chlorophyll a, conductivity, dissolved Oxygen (DO), and temperature was collected from 2010 to 2013. Secchi depth data are available from 2012 and 2015. Metagenome samples were collected in 2015.
