## Supplementary material for "Depth-discrete metagenomics reveals the roles of microbes in biogeochemical cycling in the tropical freshwater Lake Tanganyika": Figure S2

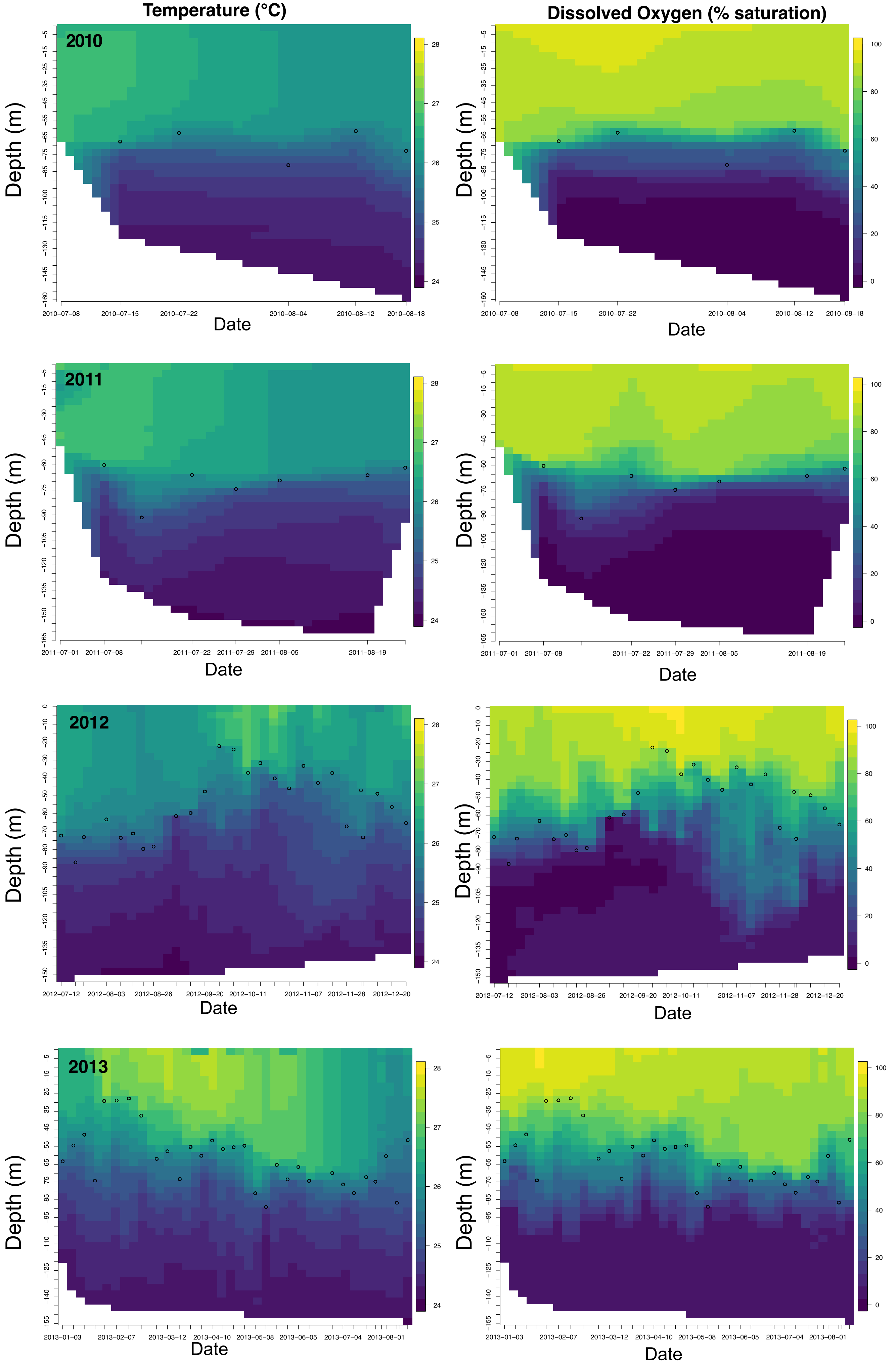


**Supplementary Figure 2.** Environmental Data Profiles from 2010-2013. Dots represent the thermocline depth at each sampling date.
