## Supplementary material for "Depth-discrete metagenomics reveals the roles of microbes in biogeochemical cycling in the tropical freshwater Lake Tanganyika": Figure S3

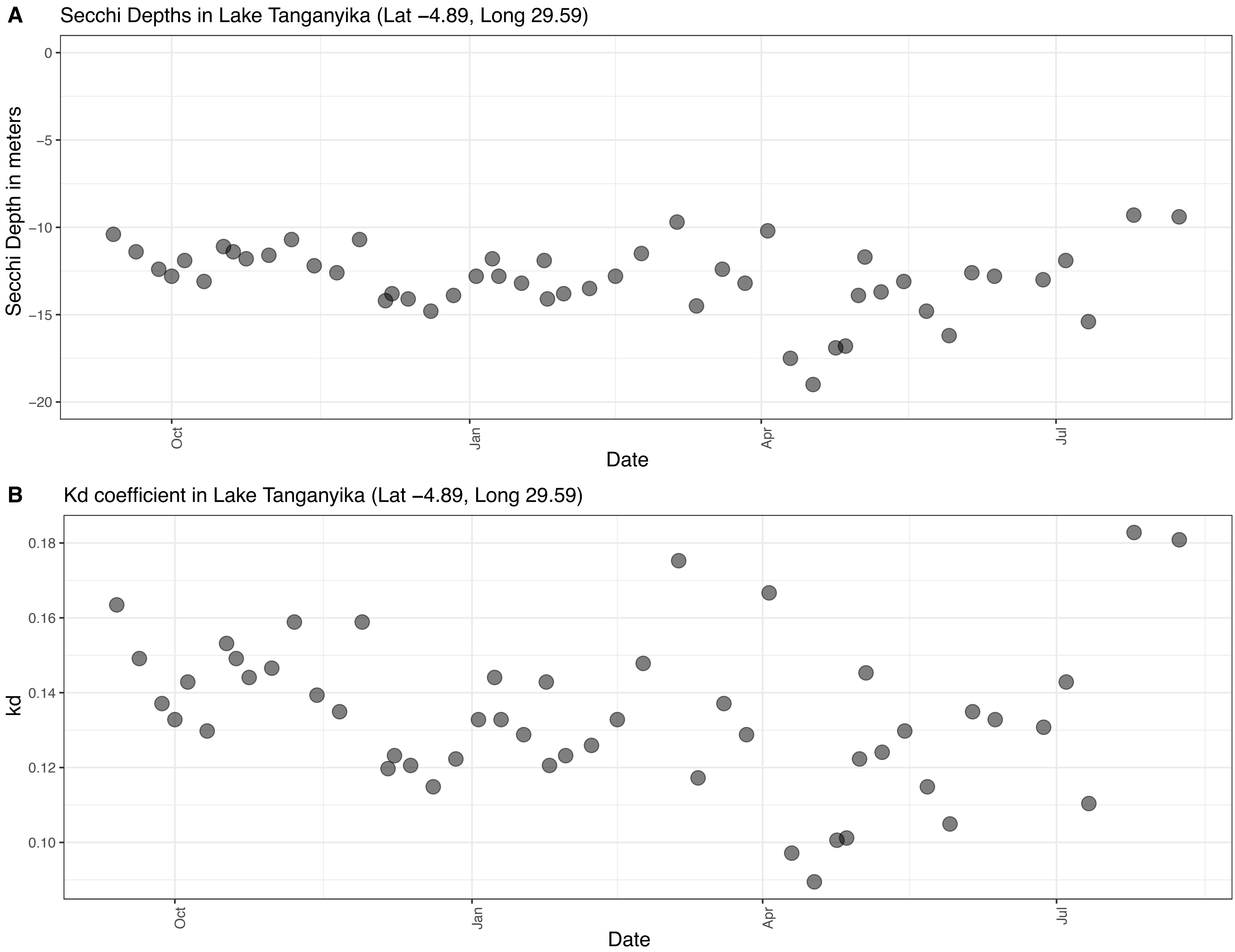


**Supplementary Figure 3.** A. Secchi depths in Lake Tanganyika. B. Kd coefficient in Lake Tanganyika, calculated by 1.7 divided by the Secchi depth in meters. Location Lat -4.89, Long 29.59 is approximately where the metagenome : KigOffshore Cast was taken, which consists of samples KigOff0, KigOff40, KigOff80 and KigOff120. Based on this trend of the Secchi depths presented in this figure, we estimate that all three metagenome KigOff40, 80, and 120 would be below the Secchi Depth.
