## Supplementary material for "Depth-discrete metagenomics reveals the roles of microbes in biogeochemical cycling in the tropical freshwater Lake Tanganyika": Figure S4

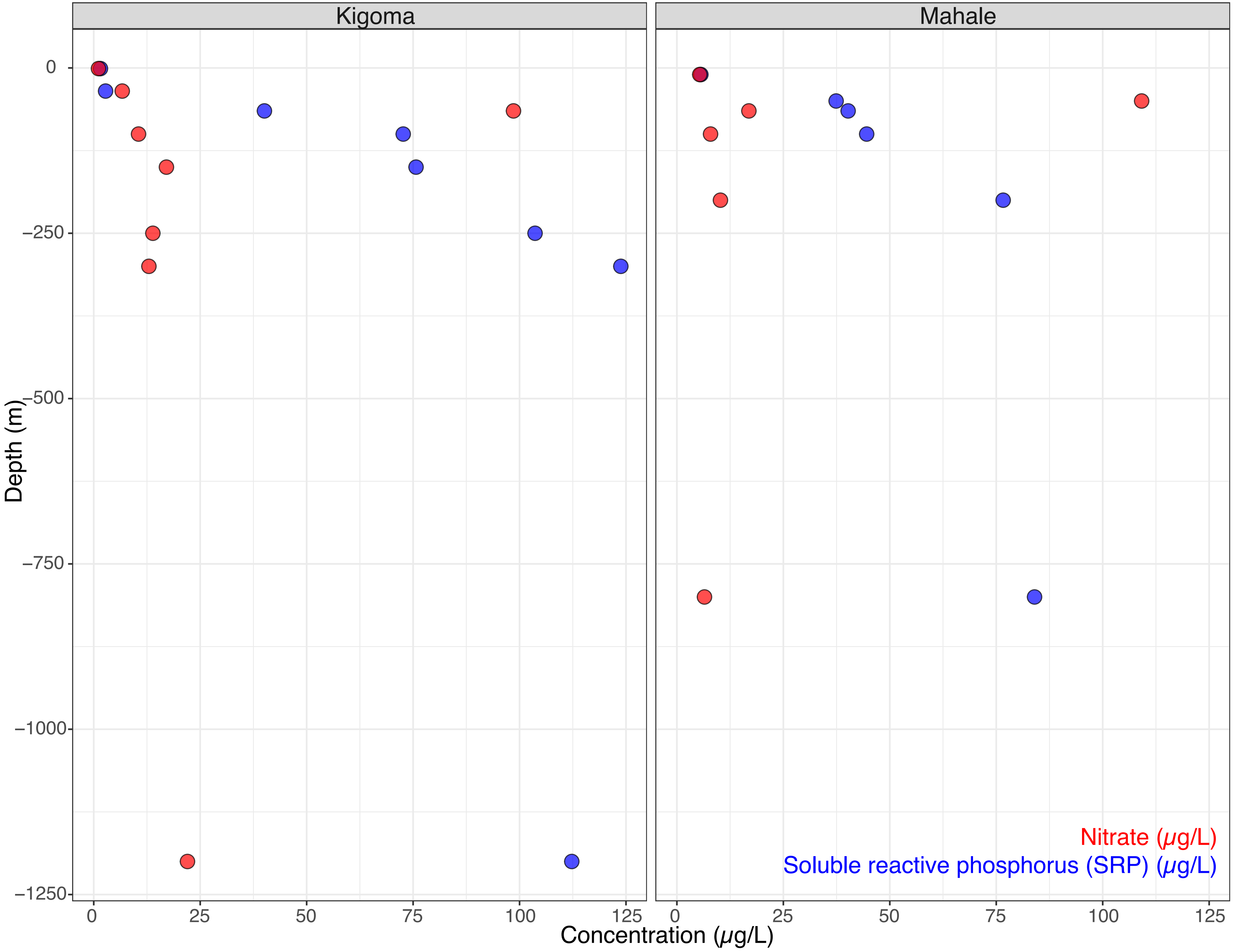


**Supplementary Figure 4.** Soluble reactive phosphorus (SRP) concentrations, which is a limnological indicator of phosphorus (blue), and nitrate concentration (red) in Lake Tanganyika. Data was collected on July 15, 2015 in Lake Tanganyika at the Kigoma and Mahale station. The peak in nitrate occurred between 50 and 65m.
