## Supplementary material for "Depth-discrete metagenomics reveals the roles of microbes in biogeochemical cycling in the tropical freshwater Lake Tanganyika": Figure S5

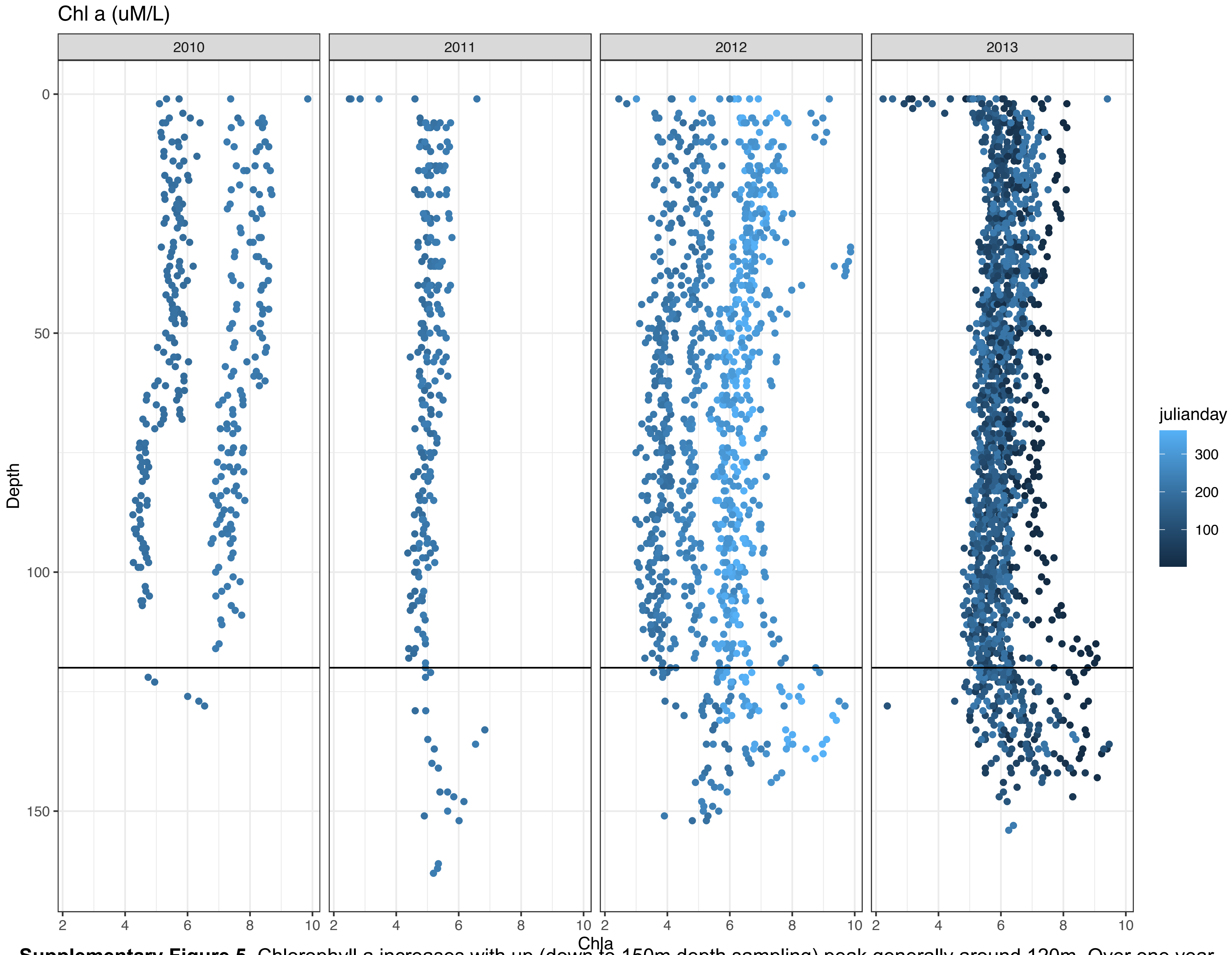


**Supplementary Figure 5.** Chlorophyll a increases with up (down to 150m depth sampling) peak generally around 120m. Over one year, chlorophyll a increases across all depth (points shift towards the right).
