## Supplementary material for "Depth-discrete metagenomics reveals the roles of microbes in biogeochemical cycling in the tropical freshwater Lake Tanganyika": Figure S6

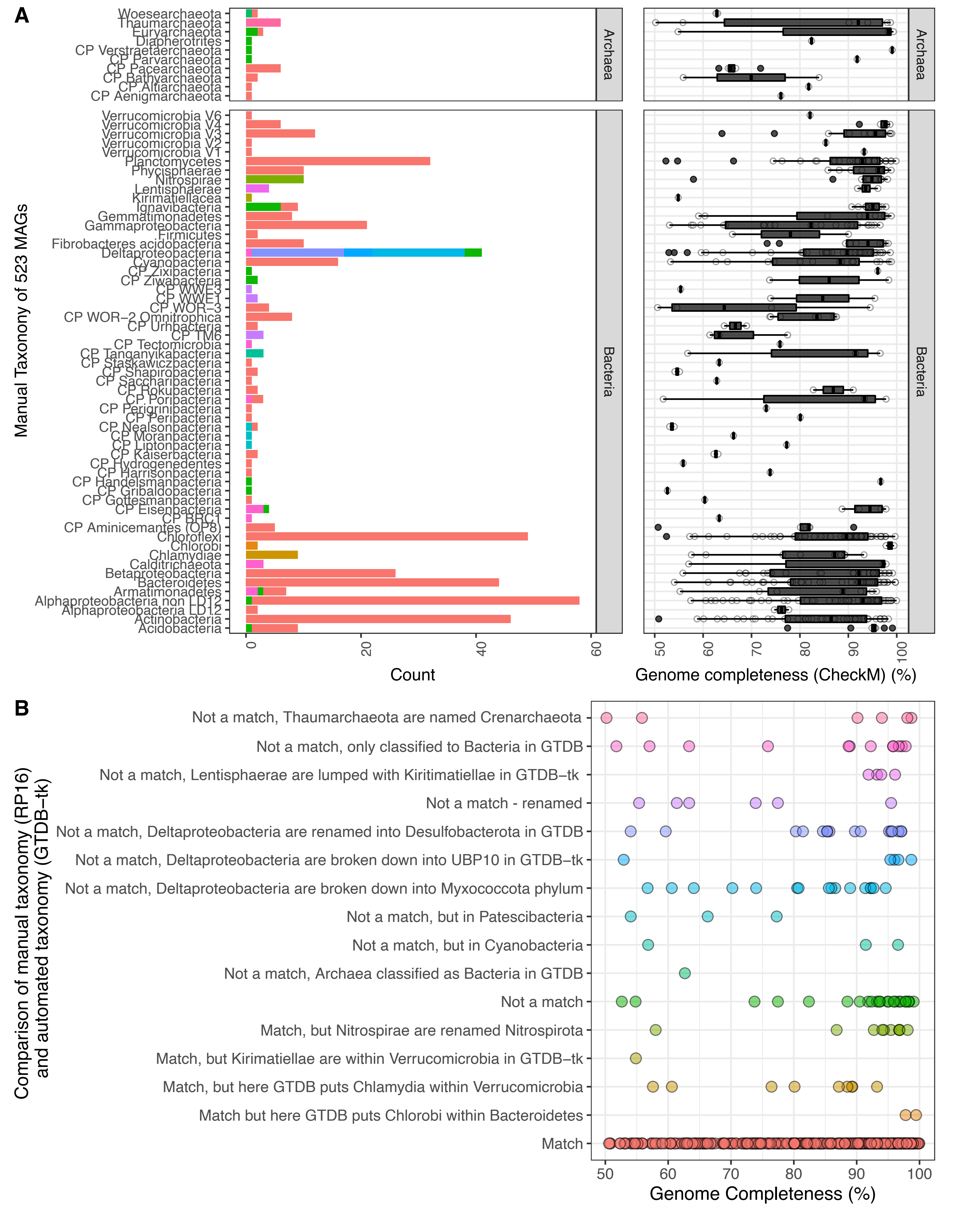


**Supplementary Figure 6.** A. Comparison of manual RP16 taxonomy versus the GTDB-tk automated taxonomy, and distribution of completeness levels on for each taxonomy group. B. Distribution of completeness values among categories. The colors in panel B is the legend for the colors in the bar plot of panel A. There were matches across the range of completeness values (“Match”). However, when there is a mismatch, it is not always because the genome is less complete.
