## Supplementary material for "Depth-discrete metagenomics reveals the roles of microbes in biogeochemical cycling in the tropical freshwater Lake Tanganyika": Figure S7

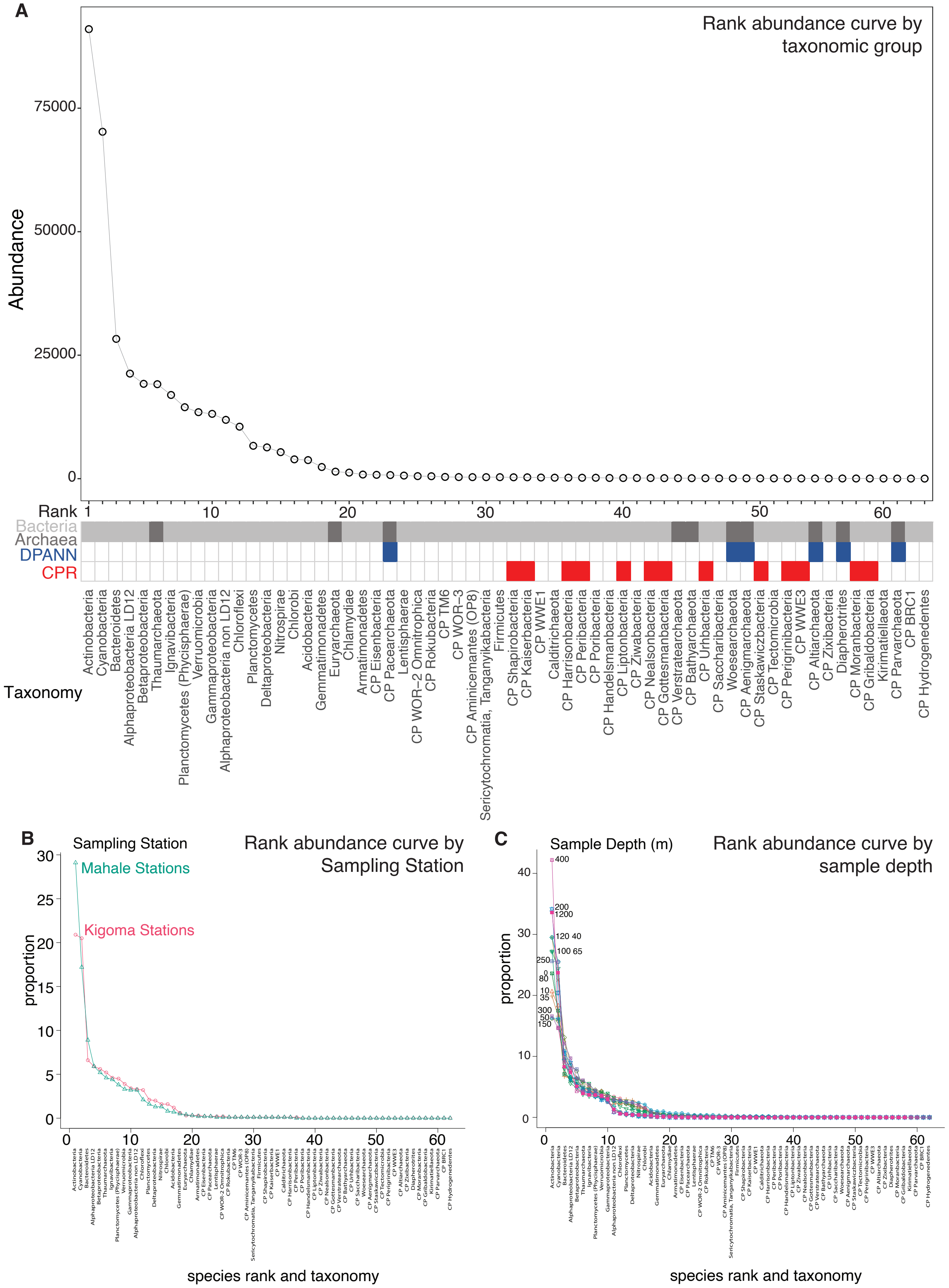


**Supplementary Figure 7.** A. Rank abundance curve overall for all samples
B. Rank Abundance curve by station C. Rank abundance curve for all samples, grouped by sample depth (meters).
