## Supplementary material for "Depth-discrete metagenomics reveals the roles of microbes in biogeochemical cycling in the tropical freshwater Lake Tanganyika": Figure S8

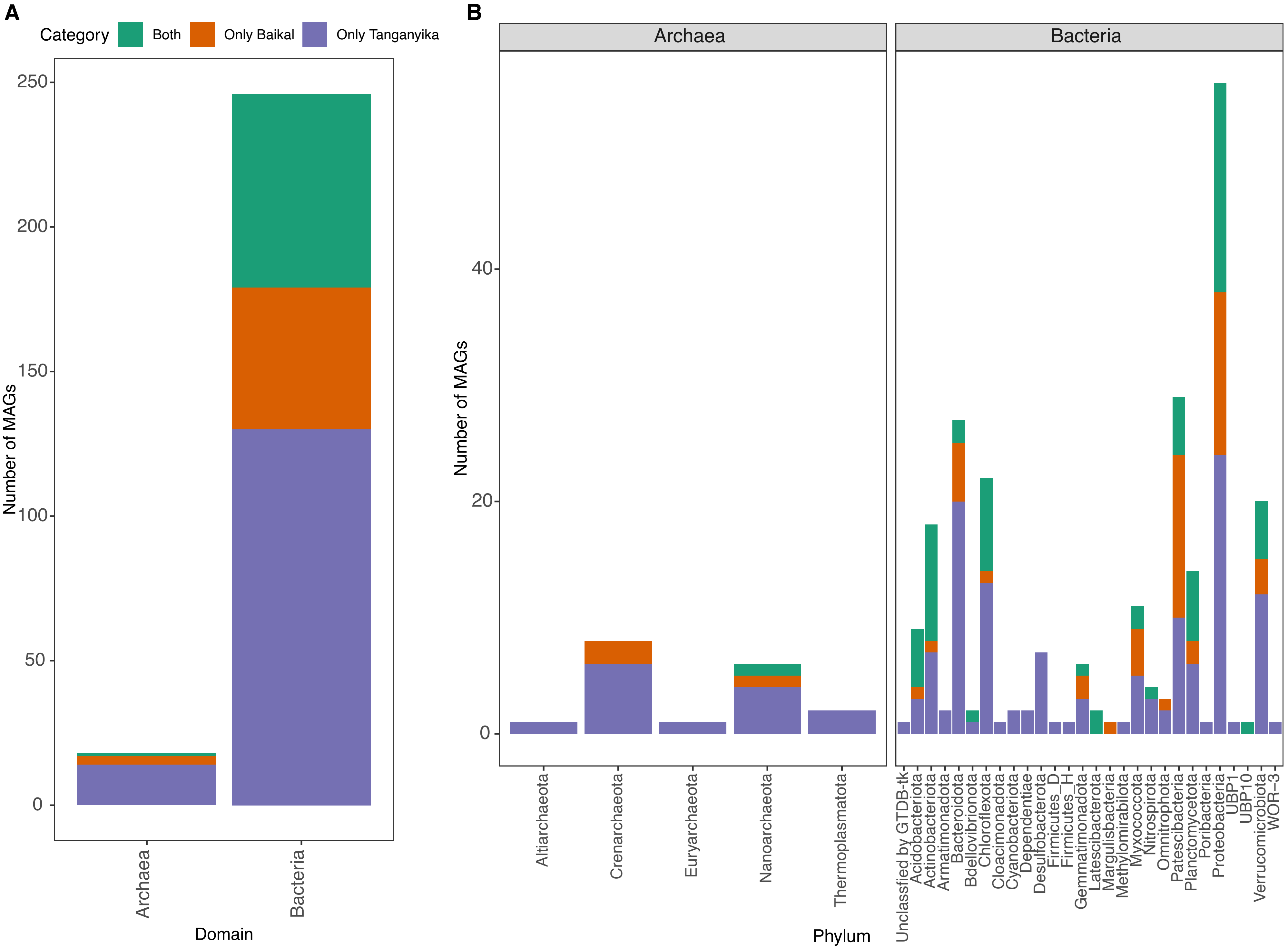


**Supplementary Figure 8.** Comparison of the taxonomic diversity between Lake Baikal and Lake Tanganyika. A. Number of MAGS shared between the two lakes and unique to either lake. B. Comparison of taxonomic diversity at the phylum-level between the two lakes.
