## Supplementary figures and images for "Depth-discrete metagenomics reveals the roles of microbes in biogeochemical cycling in the tropical freshwater Lake Tanganyika"

### Figure S9

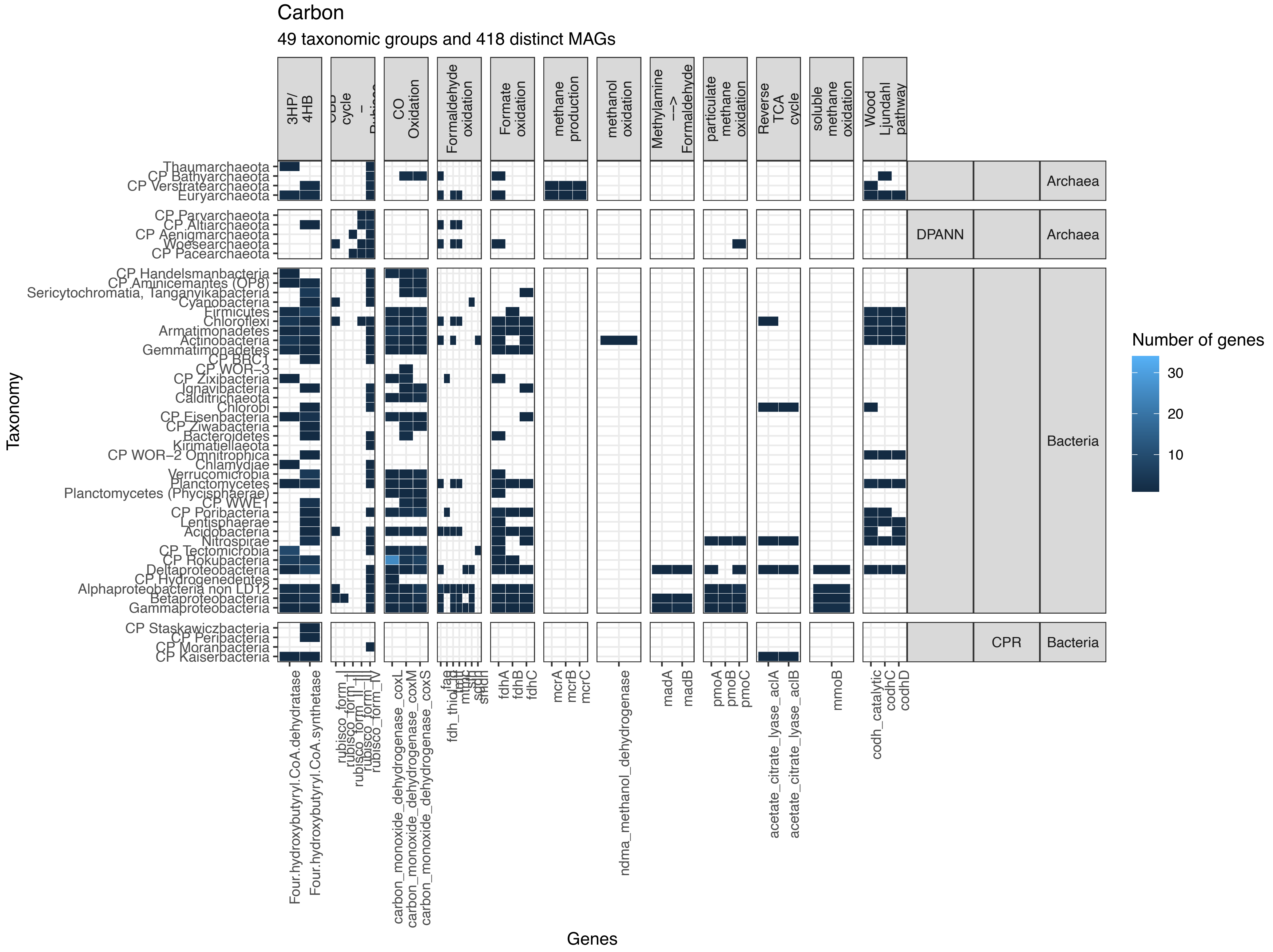


**Supplementary Figure 9.** Heatmap showing the genes involved in carbon cycling found in the MAGs.

### Figure S10

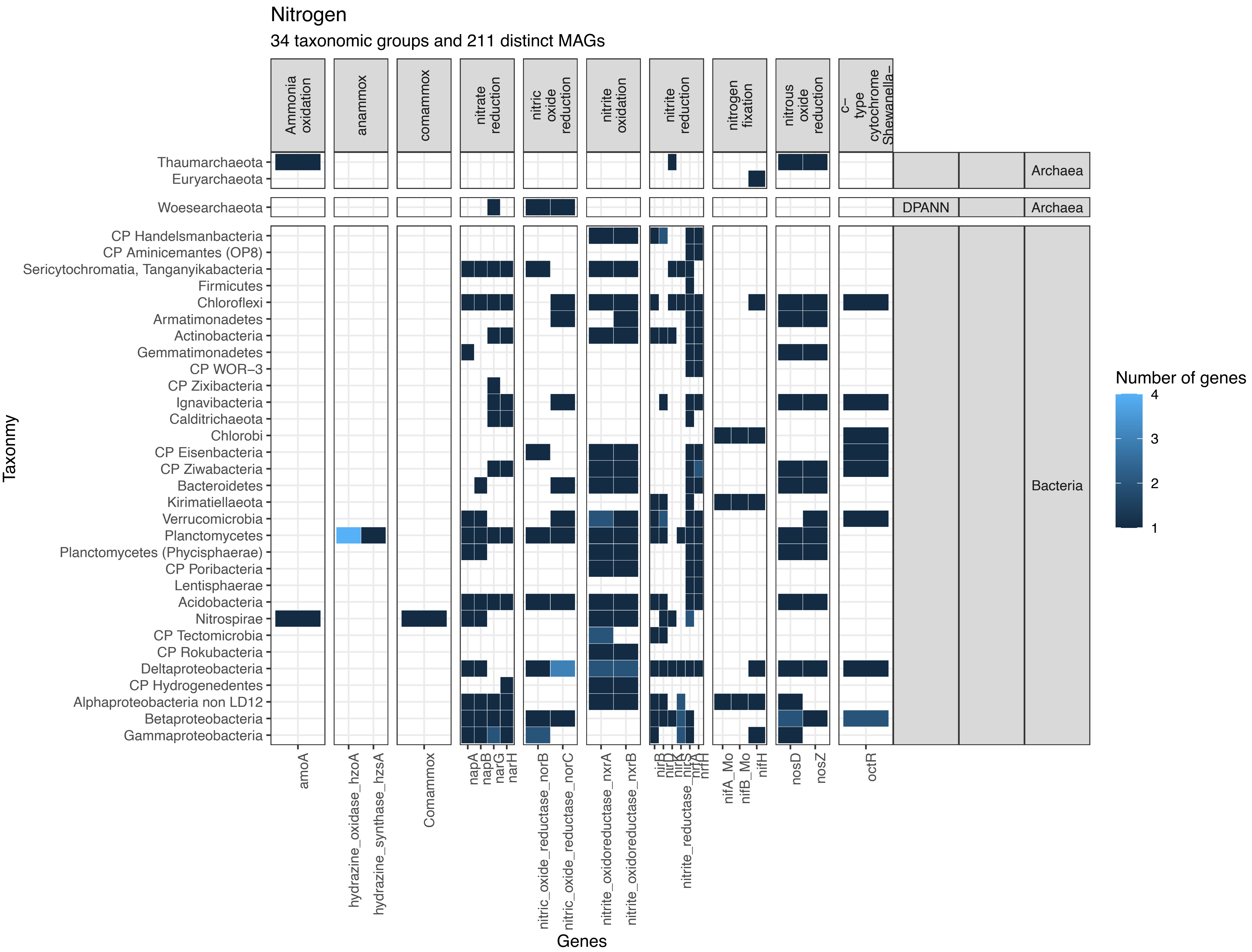


**Supplementary Figure 10.** Heatmap showing the genes involved in nitrogen cycling in the MAGs.

### Figure S11

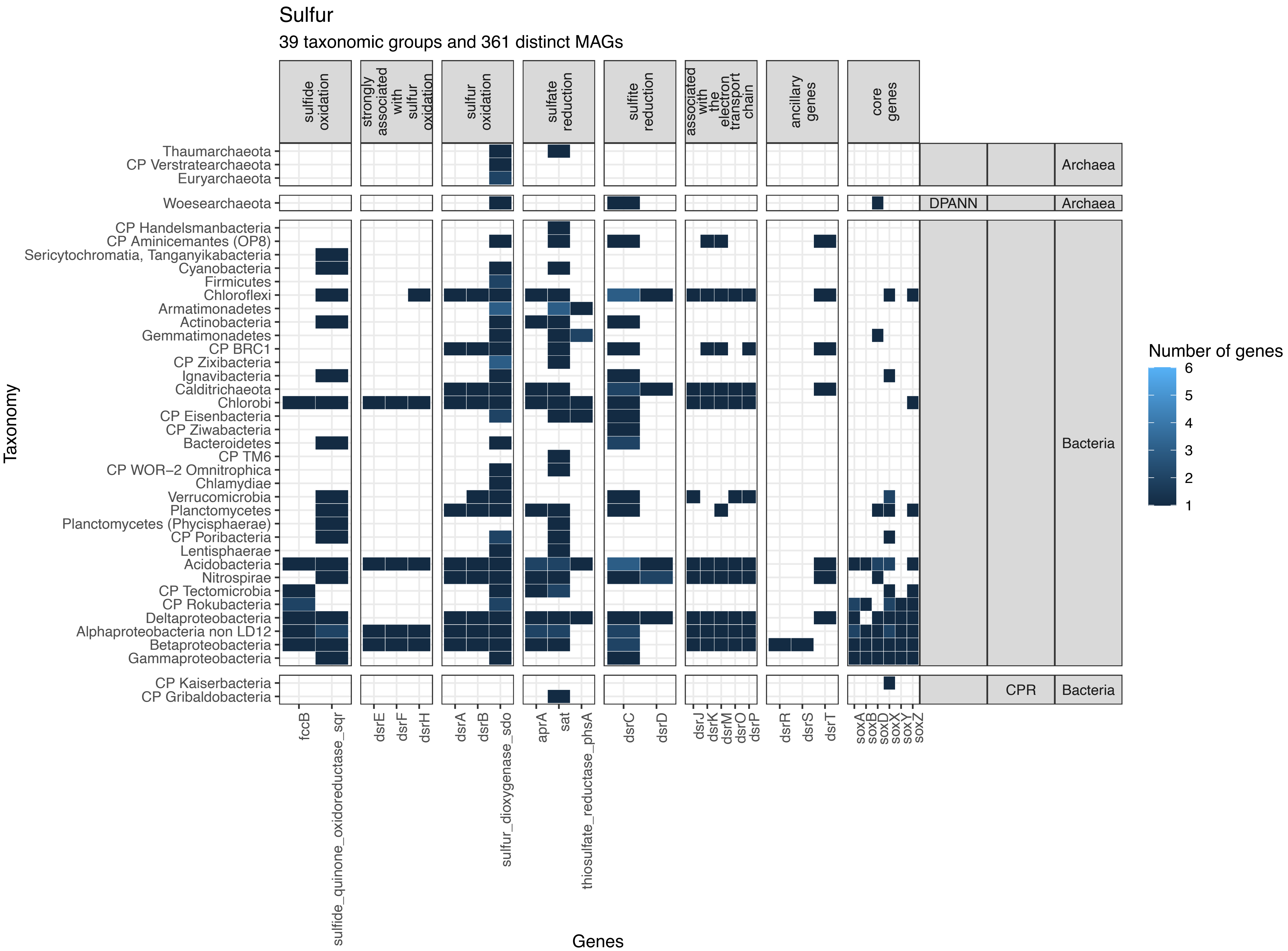


**Supplementary Figure 11.** Heatmap showing the genes involved in sulfur cycling found in the MAGs.
