## Supplementary material for "Depth-discrete metagenomics reveals the roles of microbes in biogeochemical cycling in the tropical freshwater Lake Tanganyika": Figure S12

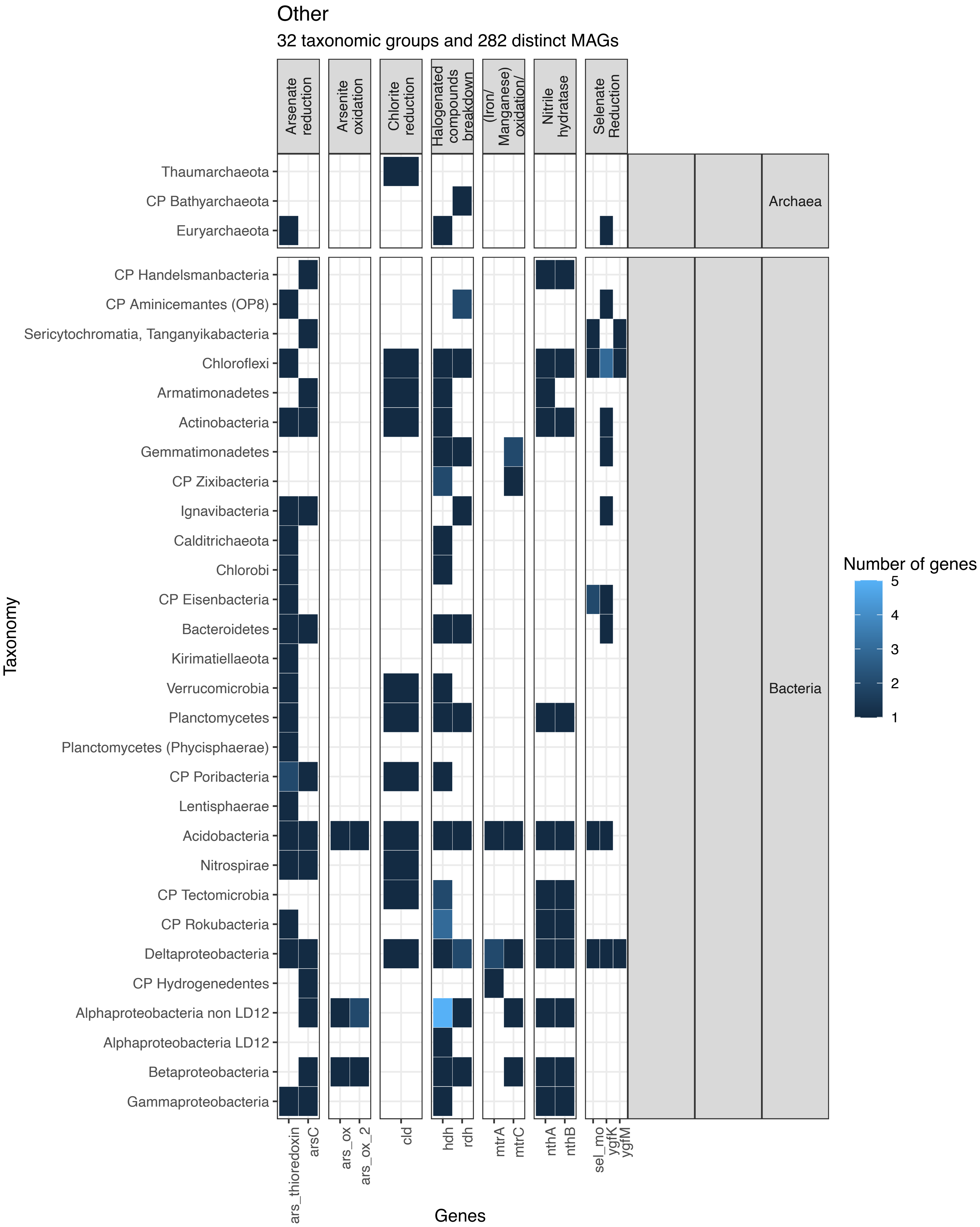


**Supplementary Figure 12.** Heatmap showing the genes involved in metal biogeochemical cycling found in the MAGs, for other types of metabolism.
