## Supplementary material for "Depth-discrete metagenomics reveals the roles of microbes in biogeochemical cycling in the tropical freshwater Lake Tanganyika": Figure S13

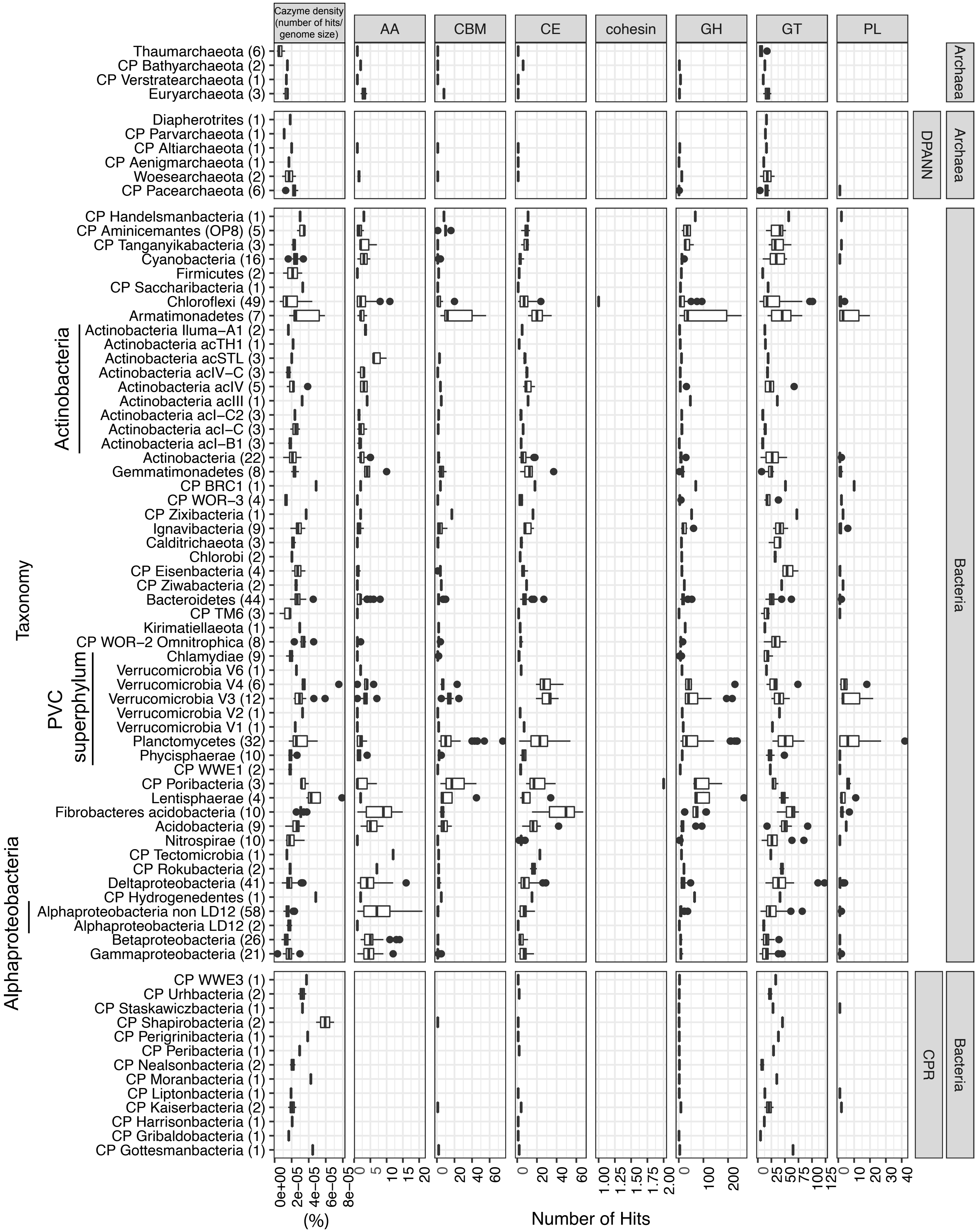


**Supplementary Figure 13.** Distribution of carbohydrate active enzymes in LT. Bar plot showing the Cazyme densities (number of cazyme hits divided by genome size in bp, in percentage), and number of hits to each CAZyme category: AA, CBM (carbohydrate binding module), CE, cohesin, GH (glycoside hydrolases), GT (glycosyl transferase), PL (polysaccharide lyase). The taxonomic groups are on the y-axis, and the number of MAGs corresponding to each group is listed in parentheses. Vertical grey bars high-light grouping of Verrucomicrobia (which are split into Orders), Alphaproteobacteria, and Actinobacteria.
