## Supplementary material for "Depth-discrete metagenomics reveals the roles of microbes in biogeochemical cycling in the tropical freshwater Lake Tanganyika": Figure S14

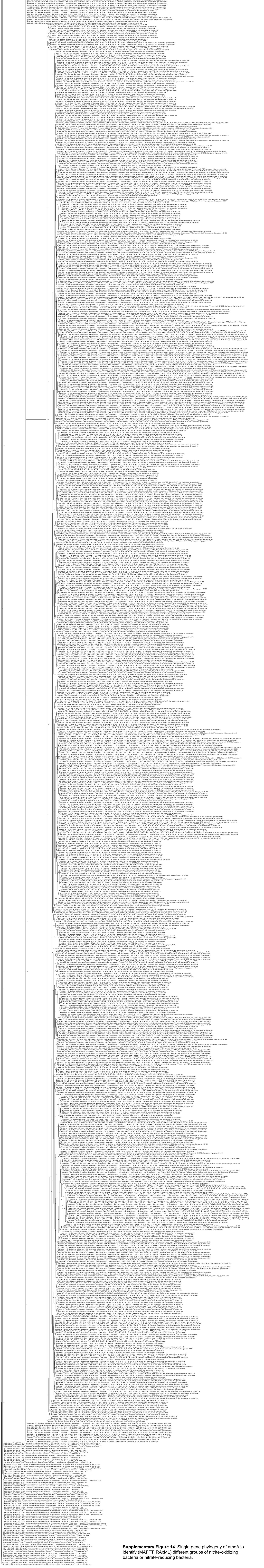

**Supplementary Figure 14.** Single-gene phylogeny of amoA to identify (MAFFT, RAxML) different groups of nitrite-oxidizing bacteria or nitrate-reducing bacteria.
