## Supplementary material for "Depth-discrete metagenomics reveals the roles of microbes in biogeochemical cycling in the tropical freshwater Lake Tanganyika": Figure S15

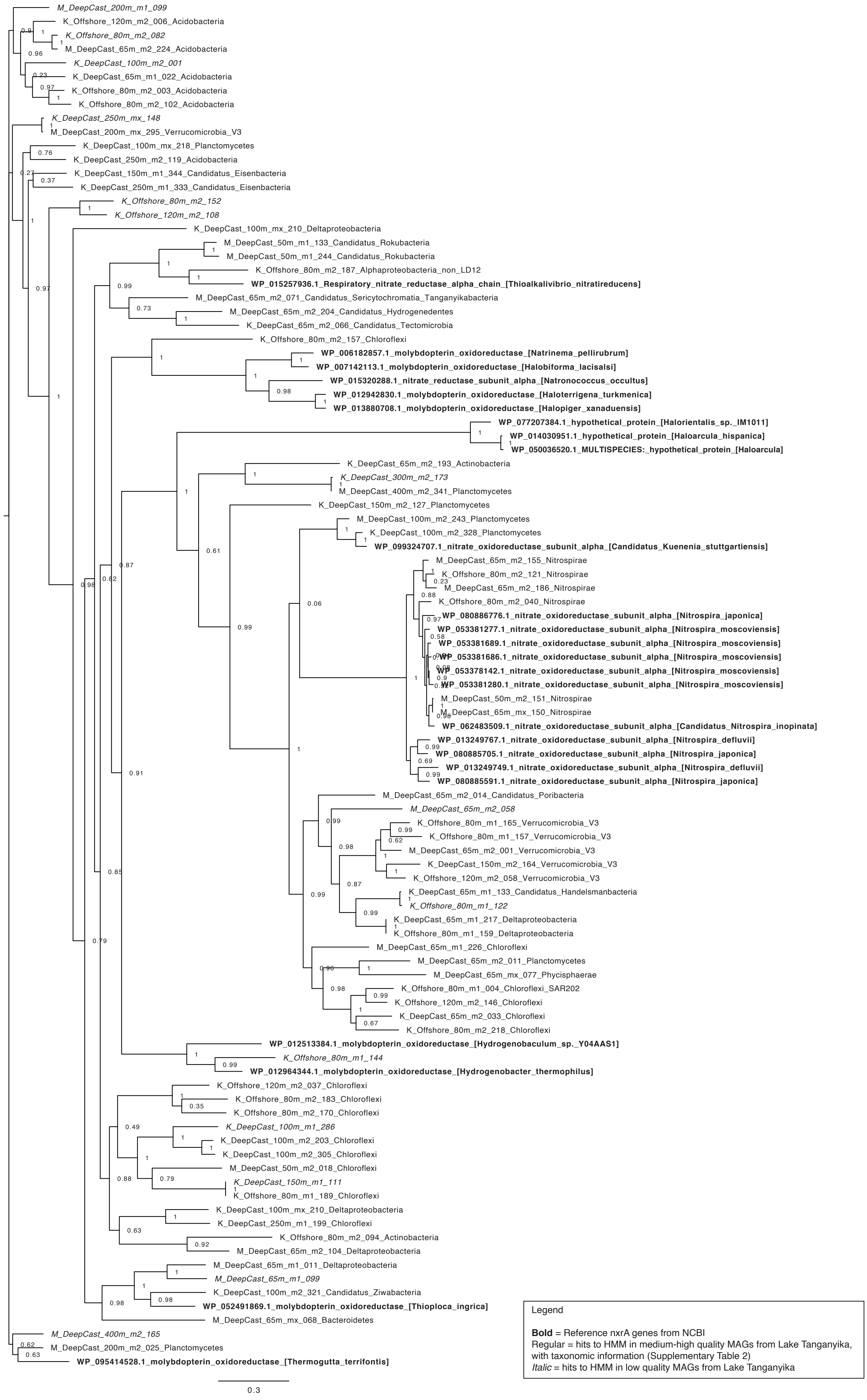


**Supplementary Figure 15.** Single-gene phylogeny of nxrA to identify (MAFFT, RAxML) different groups of nitrite-oxidizing bacteria or nitrate-reducing bacteria.
