## Supplementary material for "Depth-discrete metagenomics reveals the roles of microbes in biogeochemical cycling in the tropical freshwater Lake Tanganyika": Figure S16

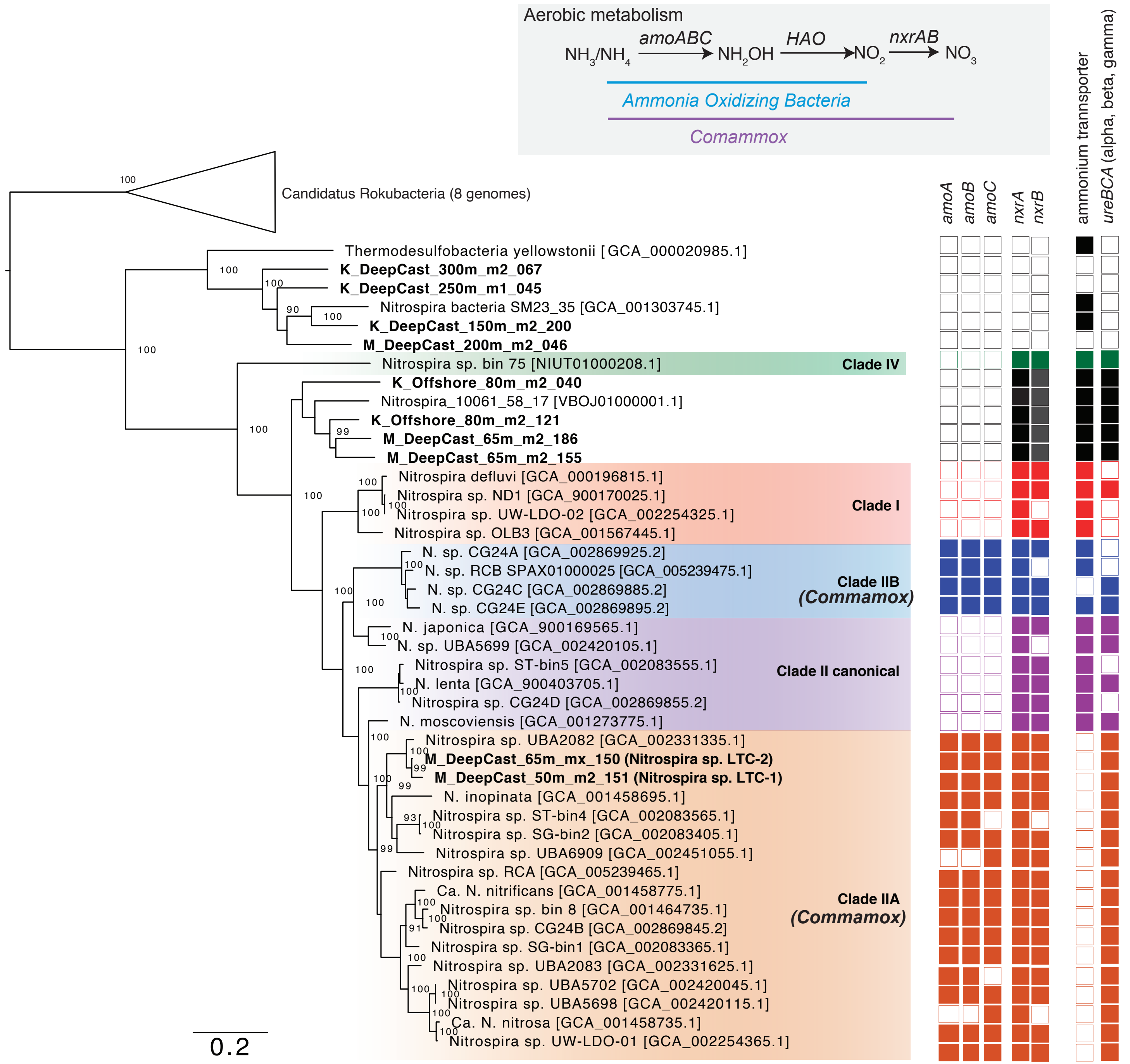


**Supplementary Figure 16.** Expanded figure showing the comammox Nitrospira from Lake Tanganyika in the context of references comammox genomes. A. Concatenated RP16 gene phylogeny of Nitrospira genomes from LT (renamed LTC-1 and LTC-2) and other environments (Accession Numbers in parentheses). B. Presence (filled boxes) and absence (empty boxes) of selected genes involved in ammonia and nitrite oxidation. The presence of comammox is based on the presence of both amo and nxr genes. ammonium transporters and urea utilization.
